# The avian colourscape is disproportionately threatened by species extinctions

**DOI:** 10.64898/2026.06.22.733718

**Authors:** Robert X. MacDonald, Kathryn Harris, Yichen He, Emma C. Hughes, Eleftherios Ioannou, Tamora D. James, Michael D. Jardine, Christopher J. A Moody, Lara O. Nouri, Zoë K. Varley, Gavin H. Thomas, Christopher R. Cooney

**Affiliations:** Ecology and Evolutionary Biology, School of Biosciences, University of Sheffield; Alfred Denny Building, University of Sheffield, Western Bank, Sheffield S10 2TN, UK; Research Software Engineering, School of Computer Science, University of Sheffield, Sheffield S1 4DP, UK; Bird Group, Natural History Museum at Tring, Akeman Street, Tring, Hertfordshire HP23 6AP, UK

## Abstract

The impact of projected extinctions on global animal colour diversity remains unknown. Combining citizen science with self-supervised deep learning, we built novel representations of bird plumage colour patterning based on >125,000 museum specimen images covering 9,143 species. We demonstrate that losing currently threatened bird species will drive a disproportionate reduction in avian plumage diversity, with the most severe losses occurring in tropical and subtropical regions. Furthermore, while humans generally find non-typical plumage phenotypes more aesthetically attractive, threatened species are unexpectedly deemed less visually appealing despite their comparatively unusual plumages. Overall, our results highlight severe, imminent threats to the existing avian colourscape and raise critical questions about the future of animal colour diversity in a changing world.

## Main text

Ongoing environmental change is altering biodiversity patterns worldwide, with widespread population declines threatening substantial species loss on regional and global scales. For certain axes of biodiversity, such as species’ phylogenetic and functional traits, it is already known that projected species extinctions are non-random and would cause disproportionately severe reductions in diversity relative to baseline expectations (*1–3*). For other aspects of biodiversity, however, we not only lack insight into the likely impact of future extinctions but also a detailed understanding of the existing diversity we stand to lose. Variation in colour patterning across organisms, particularly among animal species, is one of the most striking and immediately tangible aspects of global biodiversity, with colourful organisms and visually diverse ecosystems being highly valued by humans for their aesthetic properties (*4–7*). Yet, despite decades of study and widespread popular interest in animal colouration (*8*, *9*), we have surprisingly limited insight into the scale of threats facing animal colour pattern diversity and thus the future of this highly-valued component of global biodiversity (*10*).

One reason for comparatively limited progress in this area is the formidable empirical challenge associated with effectively quantifying organismal colouration. Animal colour patterns can vary in intricate ways and adequately capturing this variation ideally requires high-resolution information (e.g. standardised image data) and multi-dimensional analysis techniques capable of effectively representing complex spectral and spatial information. Because of these challenges, many broad-scale studies of animal colouration have focused on particular aspects of colour pattern diversity [e.g. colourfulness, (*11*, *12*)], at the expense of more complete quantitative descriptions of species’ colour pattern phenotypes. Here, we addressed this longstanding empirical challenge by combining calibrated digital imaging of museum specimens for a dense sample of bird species with citizen science and machine learning approaches to provide a comprehensive characterisation of plumage diversity for the world’s birds. We used these novel representations of avian plumage colour pattern trait space — the extant avian ‘colourscape’ — to explicitly quantify the impact of projected species extinctions on bird plumage diversity and the overall alignment between plumage diversity, species’ visual aesthetic value and extinction risk across birds.

### Characterising avian plumage diversity

Our approach to characterising avian plumage colour pattern diversity is based on a dataset of high-resolution human visible light images, comprising >125,000 images of >41,000 museum bird specimens. Where possible, three specimens of each sex for each species were imaged, with each specimen photographed from three viewing angles (dorsal, lateral, ventral) to capture whole-body plumage colouration (see Materials and Methods). Overall, we obtained images for 9,143 species across the avian radiation, sampling an average of 2.4 male and 2.1 female (and 0.8 unsexed) individuals across all taxa. We then employed two different approaches for quantifying specimen colour pattern phenotype from these images.

First, we designed an online citizen science project (http://www.projectplumage.org/) with bespoke image annotation protocols to collect information on the locations of 10 discrete homologous plumage patches for a diverse sample of specimen images. After screening, plumage patch coordinates for 24,947 specimen images were used to train and validate deep learning models (*13*, *14*) capable of efficiently predicting the location of plumage patches across the full ∼125K image dataset (see Materials and Methods). Predicted coordinates were used to extract calibrated reflectance measurements for all patches on each specimen, which were then mapped into a perceptually uniform human vision colour space (CIE LAB). This approach resulted in a 30-dimension (D) trait space capturing patch-level plumage variation across specimen images.

Second, we used a self-supervised deep learning approach (SimCLR) to encode information on specimen plumage patterning directly from images without any human labelling (*15*). This method allows the objective quantification of whole-specimen colour pattern variation and by using image augmentation during training, models can be forced to ignore non-focal features (e.g. shape variation) thereby maximising the contribution of specimen colour patterning to the resulting embeddings (*16*). We modified the original implementation of SimCLR based on a ResNet-50 backbone to explore the performance of more recent transformer-based architectures (ViT, DINOv3), finding that ViT achieved superior performance (see Materials and Methods). Separate models were trained on segmented specimen images for each viewing angle, and the resulting embeddings were concatenated to produce a 3072-D trait space capturing variation in whole-body plumage phenotype across specimen images.

Our representations of bird plumage patterning provide detailed quantitative insight into the nature of the extant avian colourscape. Principal component (PC) axes accounting for the most variance in both spaces primarily reflect variation in overall plumage lightness among specimens (i.e. achromatic variation), followed by dominant features of chromatic variation and more complex spatiochromatic plumage features (e.g. countershading) (Fig. S1). Individual PCs accounted for relatively small proportions of the overall colour pattern variation captured by each trait space (Fig. S2), implying high dimensional variation in avian plumage patterning. We therefore used Uniform Manifold Approximation and Projection (UMAP) to project these high-dimensional trait spaces into a lower-dimensional manifold, allowing us to visualise major features of the avian colourscape while retaining more complex, non-linear relationships among trait axes. The resulting projections (Fig. 1 and S3) emphasise the complex nature of avian plumage diversity, highlighting more densely packed central areas of trait space dominated by largely brown birds (e.g. points viii and x in Fig. 1B), as well as an array of highly distinct, less common and more unusual plumage phenotypes characterised by more extreme chromatic, achromatic and spatiochromatic features. This includes clusters of species characterised by highly saturated green, blue, red and yellow plumages (points i, ii, iv and v), wholly achromatic plumages (i.e. black and/or white; points vi and ix) and colourful plumages incorporating a high degree of chromatic (and/or achromatic) contrast (Fig. 1E). We further explored the structure and interpretability of our plumage trait spaces with a series of follow-up analyses (see Materials and Methods). Results confirmed that both spaces (i) capture substantial amounts of biologically relevant colour pattern information from our specimen images and (ii) provide perceptually meaningful representations of avian plumage diversity to a human observer (see Fig. S4 and S5) against which to assess the impact of projected species extinctions.

**Fig. 1.**
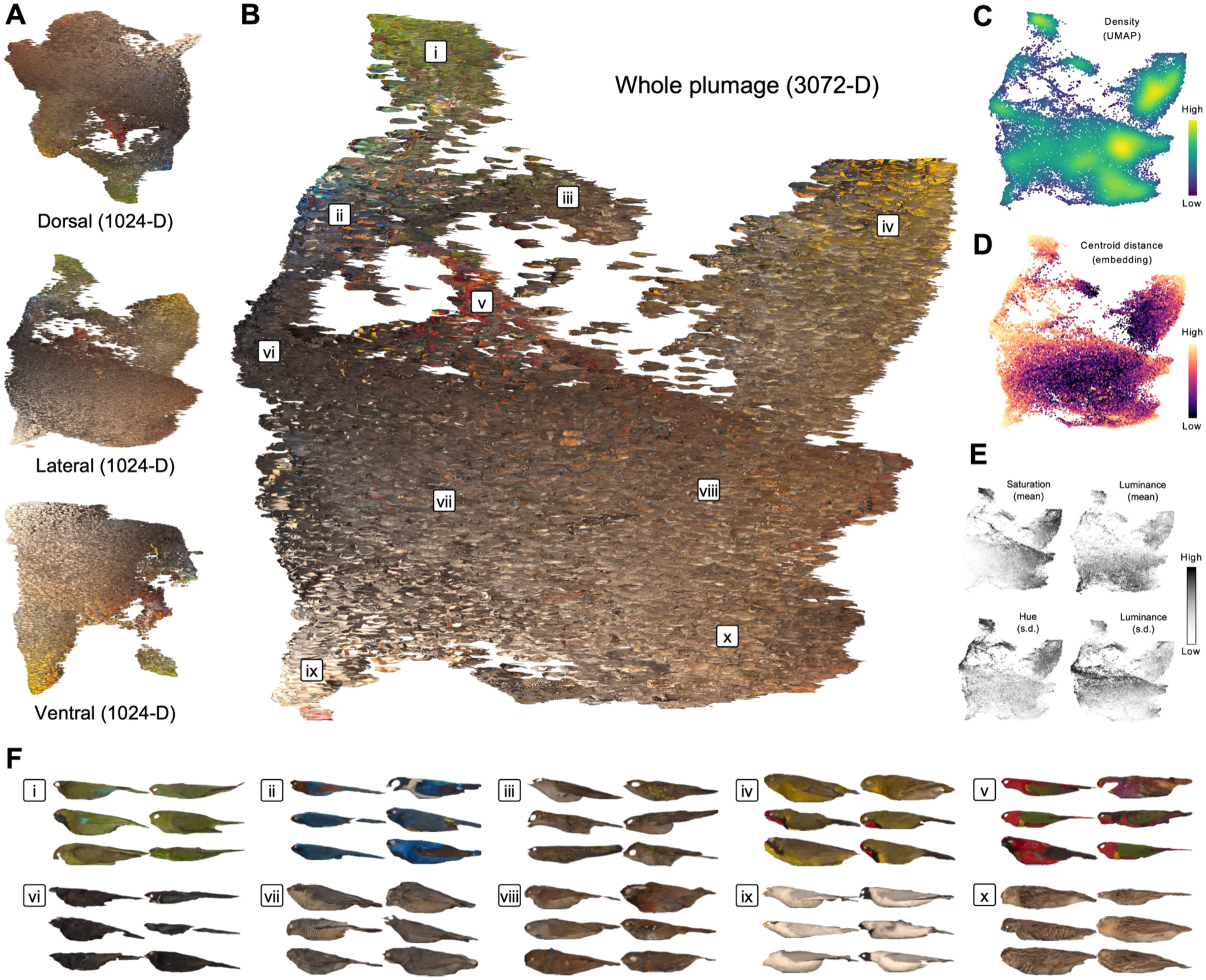
The extant avian colourscape. 2-dimensional Uniform Manifold Approximation and Projection (UMAP) visualisations of viewing angle-specific (A) and combined (B) deep learning plumage trait spaces. Note: the whole-plumage projection (B) is represented using lateral-view specimen thumbnails only. Versions of the whole-plumage projections are shown coloured according to point density in UMAP space (C), centroid distance in full 3072-D embedding trait space (D) and by selected colour pattern parameter values derived from adjacency analysis (E) (see Materials and Methods). Examples of specimens from distinct areas of the projection are also depicted (F).

### Avian plumage diversity is disproportionately threatened by extinctions

Many facets of global biodiversity are threatened by species extinctions, but animal colour pattern diversity may be at particularly severe risk. Hunting and the wildlife trade, for example, represent important anthropogenic threats to biodiversity that often directly target colourful, aesthetically attractive species (*12*, *17*, *18*). Moreover, many visually distinctive animal species inhabit geographic regions and habitat types experiencing among the highest levels of environmental change (e.g. tropical forests) (*11*), likely generating additional pressures on the extremes of animal colour pattern diversity. While there is growing appreciation that direct and indirect threats such as these may leave animal colour pattern diversity at heightened risk (*12*, *19*), for birds and most other animal taxa we still lack detailed quantitative insight into the likely impact of future extinctions on existing levels of colour pattern diversity.

To address this we tested how the global diversity of avian plumage patterning represented by our embedding spaces is impacted if bird species that are currently classified as threatened by the IUCN are progressively lost to extinction (*1–3*). To quantify colour pattern diversity, we calculated a series of trait diversity metrics suitable for high-dimensional datasets (see Materials and Methods). Our primary metric, mean distance to centroid, provides a sample-size independent measure of the average dispersion or spread of data points relative to their geometric centre, with smaller values indicating tighter clustering (lower dispersion) of observations around average trait values, indicative of phenotypic homogenisation (*20*). We performed analyses on aggregated versions of our full and dimensionality-reduced embedding spaces (*n*_spp_ = 9,143, *n*_obs_ = 16,905) as well as male- and female-only subsets (*n*_spp_ = 8,427 and 8,045, respectively), using null simulations to assess the significance of observed changes in plumage diversity relative to null models of random species loss (see Materials and Methods).

Our primary finding is that the progressive loss of currently threatened bird species causes global avian colour pattern diversity to decline significantly more severely than expected given the projected number of extinctions (Fig. 2A; Table S2). For instance, the loss of critically endangered species alone (*n* = 91) from our dataset results in a larger reduction in mean centroid distance than expected by chance, implying that losing even the most highly endangered bird species alone will cause disproportionate homogenisation of avian plumage diversity. This pattern continues as species assigned to other threat categories are further removed (Fig. 2A), with similar results for both embedding spaces and when using alternative trait diversity metrics (Fig. S6; Tables S3-5). Studying sexes separately shows that the loss of male and female plumage diversity both contribute to the overall effect, but that threatened species loss will result in relatively more severe homogenisation of female plumage diversity than male plumage diversity (Fig. 2A; Tables S2-5).

**Fig. 2.**
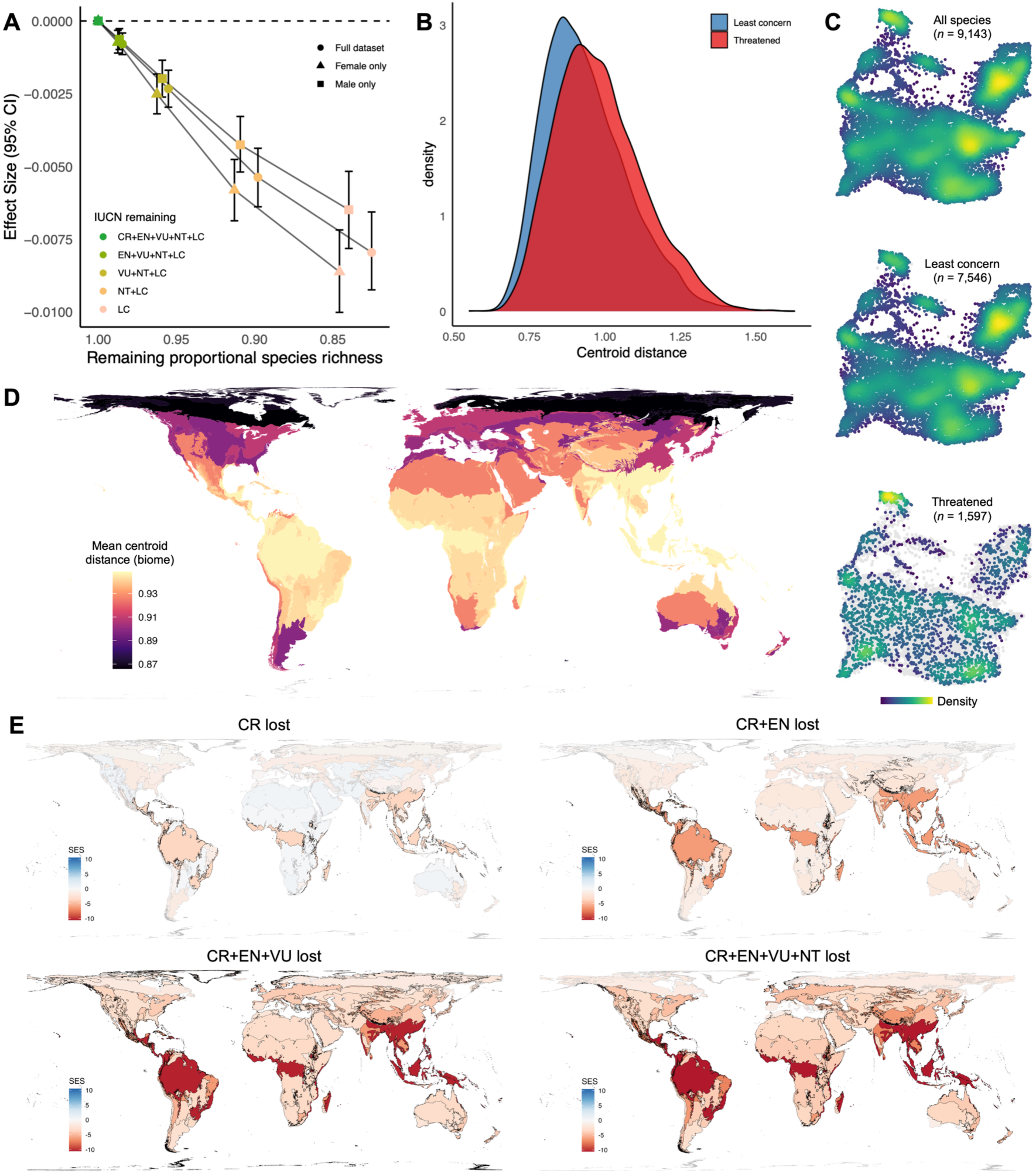
Disproportionate reductions of avian plumage diversity following threatened species loss. (A) Changes in global mean plumage centroid distance value (deep learning trait space) following sequential removal of species assigned to IUCN threat categories. Error bars that do not overlap zero (dashed line) indicates a significant (i.e. disproportionate) reduction in plumage diversity relative to null expectations. (B) Density of species’ plumage centroid distance scores, separated by threat status. (C) 2-D UMAP projections coloured by point density for different data subsets. (D) Biome-specific mean plumage centroid distance scores (*n* = 14). (E) Standardised effect sizes (SES) for biome-specific changes in mean plumage centroid distance values with the sequential removal of threatened species.

Analysing variation in centroid distance scores at the species-level shows that these assemblage-level patterns emerge because threatened and near-threatened bird species are generally distributed more towards the peripheries of plumage trait space than species of least concern (Fig. 2B-C and S6). This pattern is seemingly not driven by one or a few taxonomic groups (Fig. S7) and statistical associations between species’ threat status and plumage centroid distance scores are significant even with phylogenetic correction (Table S6). Overall, this implies a widespread association between plumage distinctiveness and extinction risk across the avian radiation. We further examined the generality of this association by re-running our simulated extinction analyses separately for major avian taxonomic groups (>30 sampled species; see Materials and Methods). Despite smaller sample sizes and reduced statistical power, overall we found that 19 (∼60%) of the 32 focal taxa showed evidence of significant reductions in plumage diversity as threatened species are lost (Table S7), confirming that these effects apply to many sub-radiations within birds as well as avian plumage diversity as a whole (Fig. S6). Examples of taxa exhibiting the strongest proportional reductions in plumage diversity after threatened species loss based on our data include the Gruiformes (cranes and allies), Trogoniformes (trogons and quetzals), Meliphagoidea (lyrebirds, bowerbirds, babblers and allies) and Galliformes (landfowl), emphasising the broad nature of the effect.

### Threats are geographically widespread but most severe in tropical regions

To evaluate how impacts are distributed geographically, we examined how extinction-driven declines in plumage diversity vary among distinct avian assemblages occupying major geographically non-overlapping terrestrial biome types (see Materials and Methods). First, our data shows that tropical and subtropical biomes (e.g. tropical rainforests, mangroves, grasslands, savannas and shrublands) currently harbour the most phenotypically diverse avian assemblages in terms of plumage colour patterning (Fig. 2D), mirroring related latitudinal trends in species’ colourfulness (*11*). Second, our data shows that overall these same tropical and subtropical regions also suffer the most proportionally severe reductions in assemblage plumage diversity following threatened species loss (Fig. 2E, Fig. S8). By contrast, losing threatened bird species from high latitude assemblages, including those found in tundra and boreal forests, has generally weaker impacts on assemblage colour pattern diversity (Fig. 2E, Fig. S8), with similar conclusions across alternative trait space and data subset combinations (Table S8). Together these results indicate that the most severe and imminent threats to avian colour pattern diversity are faced by tropical and subtropical avian assemblages. We note, however, that under an extreme scenario where all threatened and near-threatened bird species are lost, leaving only species of current least concern, disproportionate reductions in avian plumage diversity are expected in 12 (86%) of the 14 biome-specific avian assemblages studied (Fig. S8, Table S8).

### Implications for the future of avian plumage diversity

If threatened bird species do go extinct, what do we stand to lose? In terms of specific visual features, our data shows that with progressive species loss the avian colourscape would undergo significant directional shifts away from particular colours (e.g. ‘dark red’, p_9) and towards others (‘light brown’, p_7) (Fig. 3A). Likewise, threatened species loss would also cause the biased removal of particular types of plumage patterning (Fig. 3B), most notably phenotypes characterised by high achromatic contrast (m_dL), such as the black and white plumages of many seabird taxa (see below). More strikingly, however, our data shows that with threatened species loss the avian colourscape would experience widespread reductions of plumage feature diversity, indicated by smaller trait variances after species loss, caused by the non-random loss of more extreme plumage phenotypes (Table S9). This general winnowing of plumage feature diversity applies to variance both in terms of colour class proportions (Fig. 3A) and to more complex aspects of plumage colour pattern design, including colour class diversity (i.e. ‘colourfulness’; *k*, *Sc*), pattern aspect ratio (*A*) and boundary strength between colour patches (*m_dS*, *m_dL*) (Fig. 3B). Overall these results based on established hand-crafted colour pattern metrics (*21*) are highly consistent with our trait space analyses and reinforce the conclusion that the loss of threatened bird species would significantly prune the variety and composition of the existing avian colourscape across multiple dimensions of colour pattern variation..

**Fig. 3.**
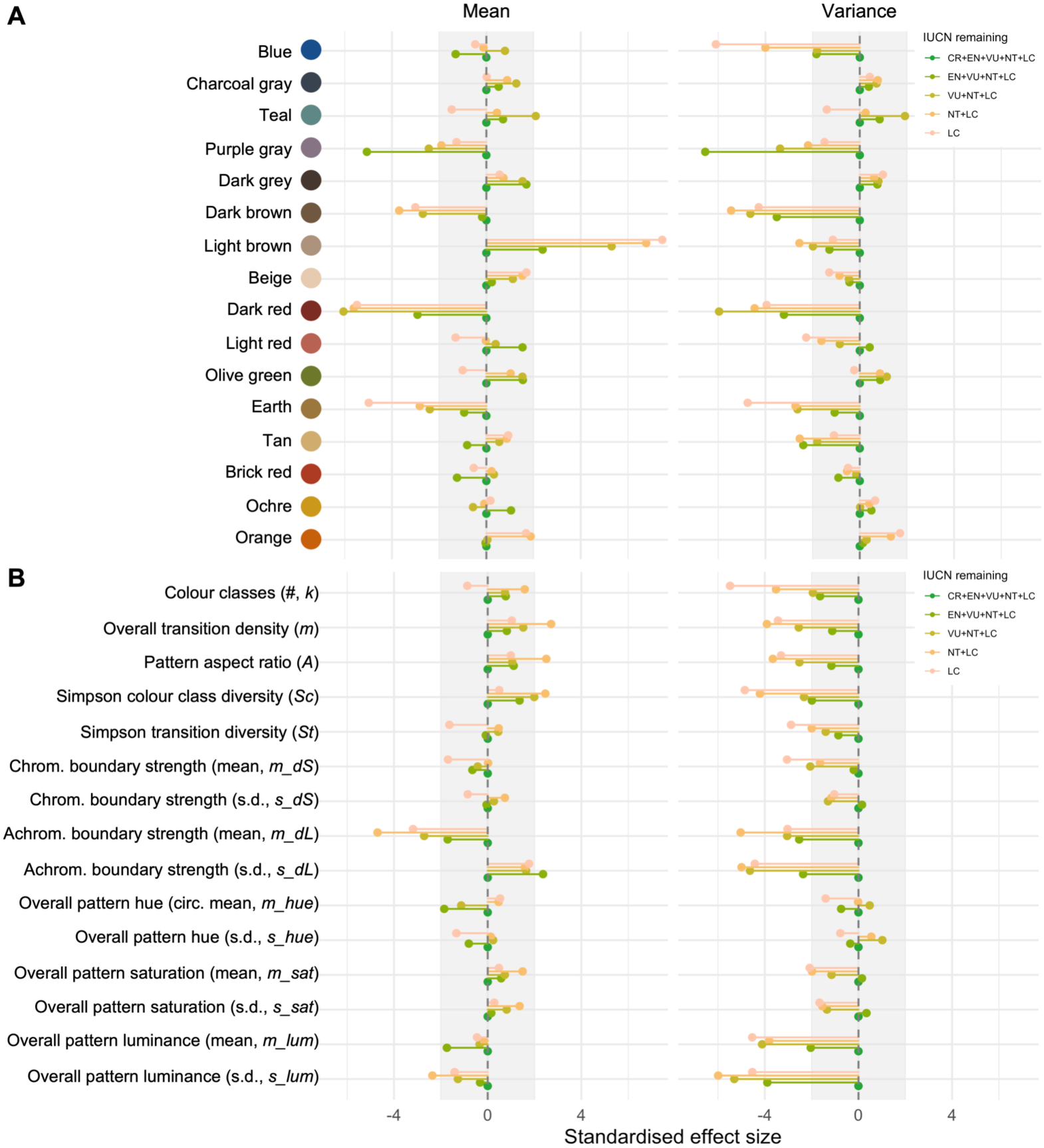
Avian colourscape changes following threatened species loss. Changes in the mean and variance of colour class proportions (A) and colour pattern parameters (B) derived from adjacency analysis following the sequential removal of threatened bird species. Points that lie outside of the shaded area (+/- 2 SES) indicate significant (i.e. disproportionate) changes relative to null expectations. Full definitions of all colour pattern parameters are provided in Tables S14 and S15. Note: colour names in A are indicative only.

So far we have considered threats to avian plumage colouration purely through the lens of biodiversity loss, but preserving animal colour pattern diversity also offers direct aesthetic benefits to humans. People generally derive considerable pleasure from viewing visually attractive organisms in nature (*4*, *22*) and for birds specifically, species with elaborate plumages containing certain colours (e.g., red, blue) are generally found to be more visually appealing (*6*). A crucial question, however, is how human-centric aesthetic preferences relate to the objective rarity of species’ visual traits (*19*). In other words, are unusually patterned bird species generally judged to be of higher visual aesthetic value? Answering this question is important both for understanding fundamental factors influencing human-nature interactions and for assessing the degree of alignment between distinct conservation-relevant facets of avian visual diversity.

We examined this by comparing our measurements of species’ plumage distinctiveness (trait space centroid distances) to an independent dataset of bird species’ visual attractiveness ratings (*23*). Attractiveness ratings provide a metric of species’ visual aesthetic value (*19*), integrating participant responses not only to species’ plumage colouration but to their overall visual appearance (*6*, *19*). Comparing the two metrics, we found a weak positive coupling between species’ plumage centroid distance scores and their corresponding visual attractiveness ratings (Table S10), implying that on average birds with more globally distinctive plumage are viewed by humans as more aesthetically attractive (Fig. 4A). However, threatened bird species are characterised by comparatively low aesthetic ratings relative to the global distinctiveness of their plumage phenotypes (Fig. 4A, Table S10), such that the relative (i.e. residual) aesthetic value of species after controlling for the overall effect of plumage distinctiveness is lower on average for threatened species than for species of least concern (Fig. 4B). Notably, we find that critically endangered species tend to be the most disproportionately unattractive relative to the global distinctiveness of their plumage phenotypes (Fig. 4, Table S10).

**Fig. 4.**
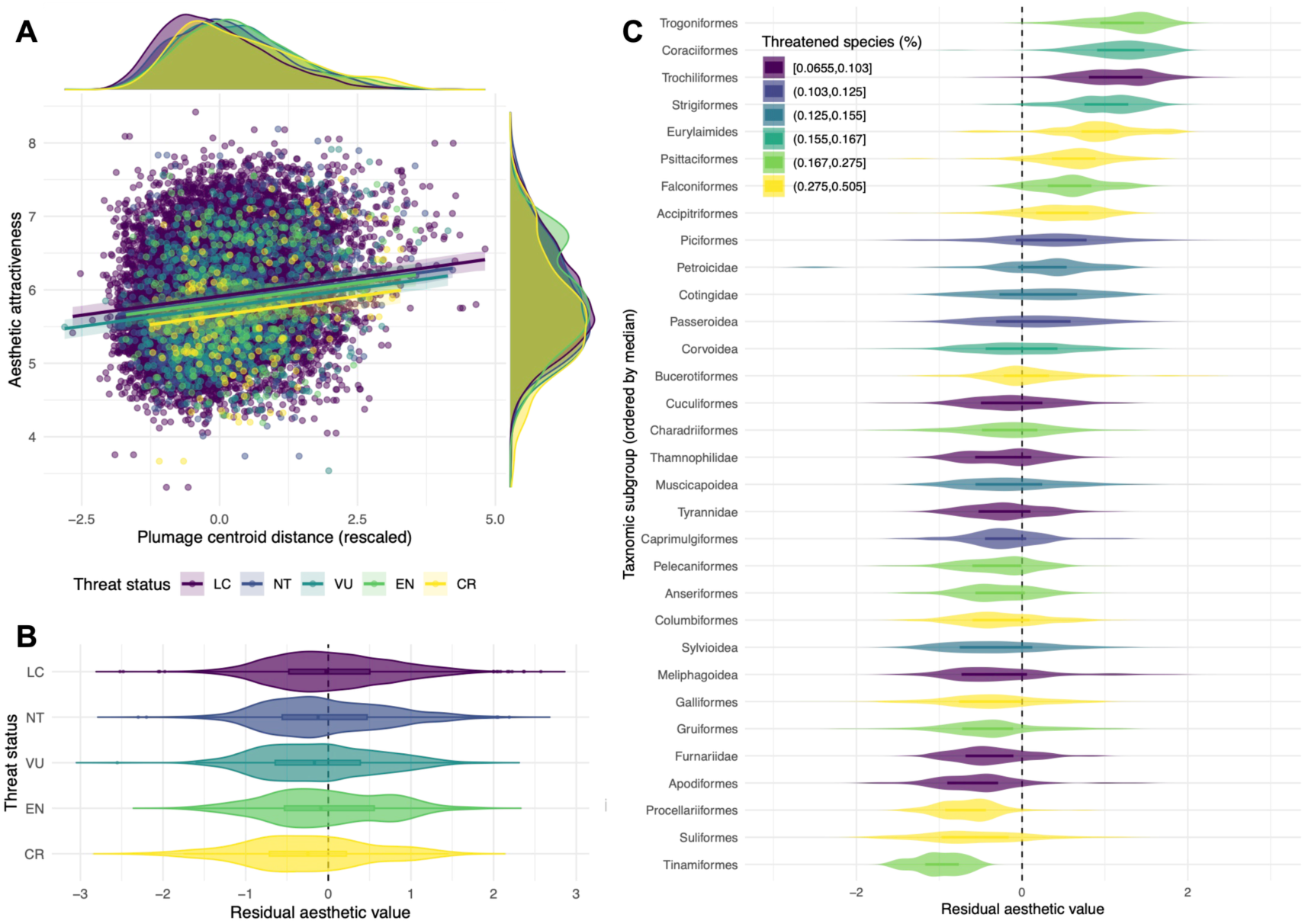
Relationships between avian plumage distinctiveness and visual aesthetic attractiveness to humans. (A) Scatterplot showing the association between species’(sex-specific) plumage centroid distance scores (deep learning embedding) and average visual aesthetic attractiveness rating, coloured by species’ threat category. Lines illustrate the fitted relationship between the variables and axis histograms indicate corresponding variable density. Also shown are the distributions of species’ residual aesthetic value scores across threat categories (B) and major taxonomic groups (C).

Separating species-specific residual aesthetic value scores by taxonomic group highlights that several avian higher taxa are consistently rated as comparatively unattractive relative to the objective global distinctiveness of their plumages (Fig. 4C). In particular, the Suliformes (gannets, boobies and allies), Procellariiformes (albatrosses, petrels and allies) and Apodiformes (swifts and allies) all contain species with predominately black and white plumages—two colours (along with yellow) that are consistently associated with lower human aesthetic value ratings (*6*). Notably, our trait spaces identify achromatic plumage phenotypes as comparatively unusual across birds as a whole (points vi and ix in Fig. 1; see also Fig. S1), providing one example of misalignment between human aesthetic preferences and plumage distinctiveness across bird species. By contrast, groups including the Trogoniformes (trogons and quetzals), Coraciiformes (kingfishers, bee-eaters and allies) and Trochiliformes (hummingbirds) are consistently rated as more visually attractive than their plumage distinctiveness would predict, possibly due to idiosyncratic plumage features in these groups (e.g. iridescence) and/or other visually appealing aspects of species’ appearance unrelated to plumage.

Overall, our data shows that while plumage distinctiveness and visual aesthetic value appear to be broadly coupled across bird species, many highly distinctive and threatened avian plumage phenotypes are not deemed particularly visually attractive by humans, and vice versa. Together, the net result of these associations is that while threatened species loss would cause outsized reductions in plumage diversity (Fig. 2), the impact on total avian visual aesthetic value would be far less severe. In fact, our analyses suggest that losing threatened bird species would result in a smaller-than-expected decrease in total avian visual aesthetic value compared to null expectations (Table S11), re-emphasising the decoupling between plumage distinctiveness and relative aesthetic value across at-risk bird species (Fig. 4).

## Conclusions

Overall, our results demonstrate a consistent and widespread link between plumage distinctiveness and extinction risk across bird species. Threatened species are at imminent risk of extinction (*24*), and so our findings highlight a severe and immediate danger to the future of the existing avian colourscape, and the direct and indirect benefits humans derive from avian plumage diversity. Across animals, more attractive species generally draw greater conservation attention, creating both opportunities and challenges for the effective allocation of conservation resources (*25–28*). By demonstrating that avian colour pattern diversity is only partially aligned with common perceptions of bird species’ visual aesthetic value, our study raises important questions regarding how both components of avian visual diversity can be optimally conserved. Addressing such questions is challenging given the range of other biodiversity axes also under threat in birds and other taxa (*1*), and moving forward it will be crucial to understand how species’ visual phenotypic traits are (or are not) aligned with other relevant biodiversity axes (e.g. phylogenetic and functional traits). In any case, our work shows that doing so will be vital for avoiding a more muted, visually homogeneous future. Finally, our study serves as a milestone in longstanding efforts to adequately quantify a prominent and valued component of global biodiversity (avian plumage) and highlights the scale of potential loss of animal colour diversity in the Anthropocene (*10*).

## Acknowledgements

We thank M. Adams, H. van Grouw, R. Prys-Jones and A. Bond from the Bird Group at the Natural History Museum at Tring for providing access to and expertise in the ornithology collection, O. Morton for helpful discussion and feedback, and all of the volunteers that contributed to Project Plumage. This publication uses data generated via the Zooniverse.org platform, development of which is funded by generous support, including a Global Impact Award from Google, and by a grant from the Alfred P. Sloan Foundation.

## Funding

The work was funded by a NERC ACCE2 DTP Studentship to RXM, a Leverhulme Early Career Fellowship (ECF-2018-101), NERC Independent Research Fellowship (NE/T01105X/1) and BBSRC International Partnering Awards Plus on AI for Bioscience (BB/Y513830/1) to CRC, and a European Research Council grant (615709, Project ‘ToLERates’) and Royal Society University Research Fellowship (UF120016, URF\R\180006) to GHT.

## Author contributions

Conceptualization: R.X.M., C.R.C.; Data curation: R.X.M., K.H., E.C.H., M.D.J., C.J.A.M., L.O.N., Z.K.V., G.H.T., C.R.C.; Formal analysis: R.X.M., Y.H., E.I., T.D.J., C.R.C.; Funding acquisition: R.X.M., G.H.T., C.R.C.; Investigation: R.X.M., C.R.C.; Methodology: R.X.M., Y.H., E.I., T.D.J., C.R.C.; Project administration: G.H.T., C.R.C.; Resources: G.H.T., C.R.C.; Software: Y.H., E.I., T.D.J.; Supervision: G.H.T., C.R.C.; Visualisation: R.X.M., C.R.C.; Writing – original draft: R.X.M., C.R.C.; Writing – review & editing: R.X.M., K.H., Y.H., E.I., T.D.J., Z.K.V., G.H.T, C.R.C.

## Competing interests

The authors declare they have no competing interests.

## Data, code, and materials availability

Data, code and materials are available at https://github.com/christophercooney/avian-colour-loss.

## Supplementary Materials

### This file includes

Materials and Methods Supplementary Text Figs. S1 to S11 References (*29–58*)

### Other Supplementary Material for this manuscript includes the following

Data S1 (Tables S1 to S14)

## Materials and Methods

### Specimen selection

We based our data collection on the taxonomic framework of Jetz *et al.* (*29*). We collected data on plumage colouration using study skins housed at the Natural History Museum, Tring, UK, with the aim of sampling as many of the Jetz *et al.* taxa as possible. The exception to this was that it was logistically unfeasible to photograph specimens of several ratite orders—specifically Struthioniformes (ostriches), Rheiformes (rheas) and Casuariiformes (cassowaries and emus)—given the large size and irregular shape of the skins, and so these groups are not represented in our analysis. For all other species, where possible we sampled specimens of three males and three females, taking care to select only mature individuals in breeding plumage with no obvious signs of moult. If no sexed specimens were available for a particular species, then we sampled up to three unsexed individuals instead. Based on these criteria we were able to sample specimens for 9,143 of the 9,993 taxa recognised by the Jetz *et al.* taxonomy, with a mean sampling of 2.4 male, 2.1 female and 0.8 unsexed specimens per species representing 41,901 specimens in total.

### Digital photography

Whole-specimen plumage colouration was measured using calibrated ultraviolet (UV) and visible (Vis) light photography, as described in detail in previous studies (*11*, *30*). Briefly, we used a modified Nikon D7000 digital single-lens reflex camera with a Nikon 105mm f/4.5 UV Nikkor lens combined with two Baader photographic lens filters: one permitting human visible wavelengths (400–680 nm; Baader UV/IR Cut filter / L filter) and another permitting UV wavelengths (320–380 nm; Baader U-Venus-Filter). Specimens were illuminated using two Bronocolor Pulso G 1600 J lamps (with UV filters removed) connected to a single Broncolor Scoro 1600 S Power Pack. The same camera settings were used for all photographs (1/250 sec, f/16.0, ISO 100, ‘Daylight’ white balance, RAW photo format), with the exception that a higher ISO sensitivity (2000) was used for UV images to achieve correct exposure. Each image also contained five Labsphere Spectralon Diffuse Reflectance standards of known relative reflectance (2%, 40%, 60%, 80% and 99%). All specimens were photographed through each filter (UV, Vis) from three different angles (dorsal, lateral, ventral). Here the focus of our study is on human-visible avian plumage colour diversity, so we focused our analysis on visible light images only. Therefore in total the plumage photography dataset used for this study consisted of 3 x 41,901 = 125,703 images.

### Image processing

Following established protocols (*11*, *30*), all raw (.NEF) specimens images were linearised and exported as linear TIFF files using DCRAW (https://www.dechifro.org/dcraw/). Image pixel (RGB) values were then normalised using mean pixel intensity values from the five grey standards included in each image to control for variation in lighting conditions following existing approaches (*31*). Normalised RGB pixel values were then translated into CIE XYZ colour space using custom mapping functions generated using information on the spectral sensitivities of our imaging setup and methods available in the micaToolbox (*31*). For a detailed description and validation of this mapping approach as applied to our dataset, please see ref. (*30*). Finally, CIE XYZ values were converted to CIE LAB values using the ‘convertColor’ function in R. We focused on CIE LAB colour space as it is a perceptually uniform colour space, meaning that equal distances in the space approximate equal distances in colour perception for standard human vision. Specimen pixel values in CIE LAB space were then either analysed directly (patch-based approach) or used to output segmented specimen images (deep learning-based approach).

### Plumage patch annotation

In this study we used two different approaches to characterise avian plumage diversity based on our specimen images. The first involved extracting colour measurements for 10 standard plumage patches (crown, nape, mantle, rump, tail, wing coverts, wing primaries, throat, breast, belly) on each specimen (*30*, *32*), providing a simple, straightforward characterisation of broad colour patterning across the body (*33*, *34*). We did this by first generating a dataset of human-annotated specimen images and then used this information to train a deep learning-based model capable of accurately predicting the location of plumage patches across the full specimen image dataset. Human annotations were generated via Project Plumage (http://www.projectplumage.org/), a bespoke online citizen science project with custom image-labelling protocols created using the Zooniverse Project Builder Platform (https://www.zooniverse.org/lab). After receiving appropriate guidance, users were asked to manually identify the location of relevant plumage patches (and grey standards) on specimen images. Overall, 28,937 specimen images covering 7,764 species were labelled in this way by 2,424 volunteers giving a total dataset of 86,148 annotations; on average each specimen was annotated 2.98 times by different users. After calculating the average (median) of replicate user annotations, the resulting point coordinates were manually checked by an expert (KH) and any images with inaccurate and/or incomplete point placements were excluded. This process resulted in a final dataset of 24,947 accurately annotated images (dorsal = 7,507, lateral = 7,277, ventral = 10,163) covering 7,753 species distributed across the avian radiation. This dataset was then used to train and validate deep learning key point prediction models for each view using PhenoLearn (*13*), a python-based toolkit for generating accurate annotations of digitised specimen images (*14*). The trained models were then used to predict plumage patch locations for all specimen images. The predicted coordinates were used to extract colour information (CIE LAB pixel values; see above) for each relevant location in an image by placing a circular bounding box (radius 25 pixels) around the predicted coordinate and calculating average (mean) values across these pixels, thereby providing a single measure representing the colour of each patch in each specimen image (*30*) for use in downstream analysis.

### Image segmentation

Our second approach to characterising avian colour pattern diversity is based on self-supervised machine learning (see below), requiring segmented specimen images as input. To segment specimens from the image background, we followed the approach outlined in Cooney *et al.* (*11*). Briefly, we used trained semantic segmentation models implemented in PhenoLearn and employed previously (*11*, *35*) to generate specimen segmentation masks for all images in our dataset. Segmentation masks were manually checked and edited using PhenoLearn where necessary, before being used to output segmented specimen images for downstream analysis. Finally, to remove small segmentation artefacts (e.g. stray pixels/small unconnected areas in cleaned segmentation masks), we used the Python OpenCV library to perform a connected components analysis (connectivity = 8) to identify all distinct components within a segmentation mask. Individual components accounting for less than <10% of the total original mask area were subsequently excluded.

### Avian plumage colour pattern space construction

An important component of our study involves the construction of a comprehensive avian plumage colour pattern trait space—that is, a Euclidean space in which distance between observations (e.g. specimens) reflects overall visual dissimilarity in plumage phenotype. Various approaches exist for extracting colour pattern information from digital images of biological specimens, including those focusing exclusively on colour (chromatic) variation (e.g. colour spaces) to those also integrating information on pattern geometry (*34*). For this study, we sought to generate comprehensive representations of avian plumage diversity capturing variation in both the spectral (chromatic and achromatic) and spatial (pattern) components of species’ phenotypes. We approached this in two ways that differed in terms of both input data resolution and analytical complexity.

#### Patch embeddings

This approach consisted of performing a Principal Component Analysis (PCA) on the mean patch-wise colour measurements (CIE LAB values) extracted for each specimen across the dataset (see above), mirroring the approach used by Cooney et al. (*30*). This has the effect of representing each specimen in a 30-dimension colour pattern trait space (“embedding space”) capturing broad differences in plumage pattern phenotype.

#### Deep learning embeddings

Building on recent study of swallowtail butterflies (*16*), we quantified colour pattern diversity across bird plumages directly from images using a deep learning-based framework for unsupervised contrastive learning of visual representations: SimCLR (*15*). The method is designed to provide a distance metric between input images and can facilitate the objective quantification of colour pattern variation between biological specimen images that may differ in shape and/or other non-focal visual characteristics (e.g. occluding body parts, labels etc.). The SimCLR approach is a form of metric learning, which focuses on learning relationships (i.e. embedding distances) between individual observations rather than on the separability of classes (*36*). The focus on learning relationships and distances between observations rather than classes is crucial for many biological applications, as it allows models to capture and emphasise important features across diverse datasets, including those that include rare observations or exhibit hard-to-distinguish phenotypic differences (*37*).

A key component of the SimCLR approach is the use of augmentations (i.e. modifications) of input images to facilitate self-supervised visual representation learning. In SimCLR, augmentations of the same initial image are considered as positive pairs, while augmentations of different images are considered as negative pairs, and the model learns to obtain smaller distances for positive pairs and larger distances for negative pairs (*15*). Common augmentations include cropping, rotating, grey scaling and colour jittering, and crucially the use of augmentations allows partial control over the features used by the model during representation learning (*16*). For example, by cropping and rotating input images within a positive pair, the model can be forced to ignore variation in specimen shape and position, while ensuring that other features (e.g. colour pattern) are used as discriminative features for identifying positive and negative pairs. The result is a trained model capable of producing quantitative (vector) representations of input images within an embedding space that captures the biologically relevant variation—such as colour and pattern—while remaining invariant to ‘nuisance’ factors like specimen orientation or shape.

For this study, we modified the implementation of SimCLR employed by Puissant *et al.* (*16*) based on a ResNet-50 backbone (*38*) to explore the performance of more recent transformer-based encoder architectures, specifically Vision Transformer (ViT) (*39*) and DINOv3 (*40*). We also extended the number of available augmentations for use with SimCLR by integrating the kornia library (*41*) with our SimCLR implementation. Our novel toolkit for contrastive representation learning using SimCLR can be found on GitHub (https://github.com/ioannouE/bio-colouration), along with complete documentation and a detailed description of the approach (https://bio-colouration.readthedocs.io/en/latest/). As mentioned, the choice of augmentations during model training is crucial as it determines the features to which the method will be invariant. Here we applied eight augmentations (Table S12) that were carefully chosen to allow us to study variation in colour patterning while minimising the contribution of nuisance variation between specimens specific to our dataset, including differences in shape, pose, alignment and occlusions (*16*) (Fig. S10). We note that no colour jittering or grey-scaling augmentations were applied to avoid obfuscating focal variation in specimen colouration (i.e. hue, brightness, saturation).

Separate models using ResNet, ViT and DINOv3 backbones were trained for each imaging orientation (i.e. dorsal, lateral, ventral) using the following hyperparameters: batch size 16, training epochs 200, temperature 0.1 and with standard input image resolution of 224 x 224 pixels. After training, the output of the last convolutional layer was used as the vector representation of the images for each model, since it retains more information on image features than the output of the multilinear projection head (*15*). Output dimensions for the ResNet, ViT and DINOv3 backbones were 2048, 1024 and 1024, respectively. Embedding output quality was based on the calculation of (i) silhouette scores (Euclidean and cosine) based on classes (species x sex) with ≥2 samples; (ii) variance explained by species, sex and class (species x sex); and (iii) agreement rate with corresponding human similarity judgements (see below). Based on these criteria, we deemed ViT to be the best performing backbone for our dataset as it produced vector representations generating higher variance explained by species, sex and class, and higher human-model agreement rate scores (Table S13).

### Linear and non-linear probing of colour pattern metrics

To establish the degree to which our novel embedding spaces captured variation in established descriptors of animal colour pattern design, we employed a probing approach (*42*, *43*). We used both linear (Ridge Regression) and non-linear (Random Forest) models to assess whether colour pattern parameters derived from adjacency analysis (*21*) were not only present in the representations but also linearly decodable from them. Adjacency analysis is a commonly employed approach in animal colour pattern research (*34*) and involves taking transects across colour patterns to estimate parameters that capture not only the relative areas of each patch class but also more complex metrics of colour pattern geometry and texture (*21*). For our purposes, it provides a useful suite of established and interpretable ‘hand-crafted’ colour pattern descriptors against which to compare our novel colour pattern trait spaces.

Adjacency analysis is performed on a ‘zone map’—that is, a representation of an organisms’ colour pattern in which each pixel in an image has been assigned to one of a discrete number of colour/luminance classes. To standardise the comparison of colour patterns across multiple images, it is necessary to define a universal set of colour classes to which pixels across all images in the dataset can be meaningfully assigned. To do this we followed the approach outlined by Weller *et al.* (*44*), implemented in the R package ‘recolorize’. To define a universal set of colour classes for our dataset, we used the patch dataset (see above), which consisted of 419,010 individual measurements of avian plumage patch colouration. Initial colour clustering was performed on this dataset using a histogram-based approach, with six bins on each of the CIE LAB colour space axes. These clusters were subsequently refined using the recluster function, with a distance cutoff of 30 in CIE LAB space (*44*). This process resulted in a universal set of colour categories for our dataset consisting of 16 distinct colour classes (Fig. S11, Table S14), which was used to create zone maps for each specimen image (224 x 224 resolution). Finally, we used the recolorize_adjacency function that calls functions in the R package ‘pavo’ (*45*) to calculate adjacency analysis parameters for each image in our dataset (see Table S15 for parameter descriptions).

We performed both linear and non-linear probing of colour pattern parameters against each set of trait space axes (patch and deep-learning embeddings). For each combination of focal feature and viewing angle, the dataset was partitioned into an 80% training set and a 20% hold-out testing. We then trained two distinct predictive models for each focal feature: (1) Linear probing was performed using Ridge Regression, specifically an L2-regularised regression model built using the R package ‘glmnet’ (v.4.1-8), with the optimal lambda penalty automatically tuned via cross-validation. (2) Non-linear probing was performed using Random Forest in the ‘ranger’ package (v.0.15.1) to produce a non-linear regression ensemble. Model performance was evaluated on the unseen 20% test partition by calculating the proportion of variance explained (*R*^2^). Overall, this approach allowed us to systematically quantify how well both localised colour patch data and global deep-learning embeddings represent complex, spatial plumage colour metrics.

### Human-model agreement tests

To evaluate how well our representations of avian plumage diversity align with human perception of visual similarity, we compared human similarity judgments against distance metrics derived from each of our embedding spaces. We did this using a two alternative forced choice (2AFC) test design that asks which of two images is more similar to a reference which has been used to benchmark the performance of other visual perceptual metrics (e.g. LPIPS) (*46*). For these analyses we focused on a representative sample of dorsal specimen images, which were selected using the geosketch algorithm (*47*) applied to the dorsal deep-learning embeddings to provide a geometry-preserving sample of 5,000 images (accounting for 90% of dataset variance) for subsequent testing. We used this sample to generate 2,000 unique 2AFC trials, consisting in each case of a focal “anchor” specimen and two comparison specimens.

To represent human similarity judgements, one of us (KH) then categorised specimens in each trial as the “winner” (the specimen is more visually similar to the anchor) and the “loser” (the less similar specimen). For the model representations, we calculated the Euclidean distances between trial specimens in the embedding space and categorised the model-judged “winner” as the specimen with the smallest distance to the anchor specimen, and the “loser” as the specimen with the larger distance. Overall human-model agreement rate was thus scored as the percentage of 2AFC trails in which both human and model selected the same “winner”, and we used exact binomial tests to determine if agreement rates were significantly better than random change (50%).

We also calculated the absolute difference between the two model-based distance scores within each trial to serve as a measure of the model’s confidence and/or a proxy for the “difficulty” of the discrimination task. We then modelled the relationship between human-model agreement and the magnitude of the (normalised) distance difference in embedding space using logistic regression and used this to visualise how human-model agreement rate varies with normalised distance difference.

### Trait space similarity, structure and interpretability

We explored the similarity, structure and interpretability of our plumage trait embedding spaces with a series of follow-up analyses. First, Mantel tests based on a sample of datapoints (*n* = 5,000) revealed significant structural agreement between the patch and deep learning embedding spaces (Mantel test: *r* = 0.624, *p* = 0.001) (Fig. S4A), despite the differences between the two approaches. Second, linear and non-linear probing analyses using colour pattern metrics derived from adjacency analysis (*21*) (Fig. S5 and S11, Tables S14 and S15) showed that both embedding spaces encoded substantial amounts of meaningful colour pattern information from specimen images. Specifically, the patch and deep learning embeddings explained a median of 64% (range: 4–89%) and 90% (range: 46–99%) of the variance in the adjacency metrics, respectively (Fig. S4C, Table S1). Third, human scoring of specimen similarity using two-alternative forced choice (2AFC) tests revealed that overall human-model agreement regarding specimen similarity was 69% for the patch embedding and 75% for the deep learning embedding (Fig. S4B), with both rates significantly exceeding the 50% value expected by chance (two-sided exact binomial tests: *n =* 2,000*, p* < 0.001). Further analysis also revealed that human-model similarity agreement rate varied intuitively with the apparent difficulty of the 2AFC task – agreement rate was lower when human and model were asked to choose between two visually similar specimens (i.e. the ‘correct’ answer was largely arbitrary), but much higher when one specimen was clearly more similar to the focal image than the other (Fig. S4B). Together, these results imply that both embedding spaces capture substantial amounts of biologically relevant colour pattern information from our specimen images, providing meaningful human-centric representations of avian plumage diversity.

### Colour pattern space dimensionality estimation

Following Woo *et al.* (*48*), we used parallel analysis to estimate the dimensionality of our colour pattern trait spaces. Parallel analysis reports the number of principal components with statistically significant explanatory power compared to a null distribution of parallel PCAs in which data points have been independently shuffled, and was found to be the most accurate approach among linear methods for both linearly and nonlinearly embedded simulated data (*49*). We implemented parallel analysis with the ‘PARALLEL’ function from the ‘EFAtools’ package (v.0.4.4) in R, with 1,000 simulations. For our patch embedding, parallel analysis identified 6 relevant dimensions and for our combined deep learning representation, 129 relevant dimensions were identified. Together, these reduced sets of axes accounted for ∼75% and ∼93% of the total variation in the original trait spaces, respectively. We repeated our main analyses on these parallel analysis-reduced embedding spaces in order to assess the robustness and sensitivity of our results.

### Quantifying colour pattern diversity

We use mean distance to centroid (i.e. the average Euclidean distance of all observations to the trait space centroid) (*20*) as our primary metric of avian colour pattern diversity measured from our embedding spaces. This metric, also known as ‘functional dispersion’, measures the spread of datapoints within multivariate trait space and captures variation in both the volume and density of occupied trait space (*2*, *50*). A high mean distance to centroid value indicates that the trait values of included taxa are on average highly different (‘dispersed’) from one another in trait space. A reduction in mean centroid distance following the removal of taxa is therefore indicative of trait homogenisation—i.e. the remaining taxa are more phenotypically similar to one another, clustering more closely around the average trait value. Centroid distance values are calculated on a per-observation basis (i.e. each datapoint has its own distance to centroid) and can be interpreted as a measure of how ‘typical’ or ‘average’ a given observation is, relative to the dataset. Thus in terms of our colour pattern trait spaces, observations (e.g. specimens or species) with low centroid distance scores can be interpreted as having plumage phenotypes that are relatively typical across all birds, whereas observations with high centroid distance scores are those that exhibiting comparatively untypical or distinctive plumage phenotypes. We calculated centroid distances scores using the R package ‘dispRity’ (v.1.7.0) (*51*).

While we primarily focus on centroid distance scores as our disparity metric, we also examined the sensitivity of our primary findings to the use of other trait diversity metrics. Specifically, we employed metrics suitable for high-dimensional datasets within our global simulated extinction analyses to assess the robustness of our main result. These metrics were (i) sum of variance, (ii) sum of ranges and (iii) sum of 95% quantiles across trait space axes which represent alternative approaches for quantifying multidimensional trait space diversity (*52*). We note that other commonly used volume-based metrics of trait diversity, including convex hulls and probabilistic hypervolumes, are not applicable in our case due to the high-dimensionality of our trait data—such measures are computationally intractable for trait spaces of more than ∼5 dimensions.

### Assemblage-level analyses

Following the approach of previous studies [e.g. (*1*, *2*)], we quantified the impact of threatened species extinction on avian colour pattern diversity by performing ‘simulated extinctions’. This involved sequentially removing species at decreasing IUCN threat levels from the colour pattern space and testing how observed changes in trait diversity compared to null expectations (random species removal). Information on species’ threat level was taken from the International Union for Conservation of Nature (IUCN) Red List (version 2023-1). We matched the taxonomy used by the IUCN to the Jetz *et al.* taxonomy used in data collection using the AVONET BirdLife-BirdTree crosswalk (*53*). Species in our dataset categorised by the IUCN as extinct (EX, n = 5) or extinct in the wild (EW, n = 5) were relabelled as critically endangered to reflect the recent/ongoing nature of these impacts. Taxa in our dataset classed as data deficient (DD, n = 22) were conservatively relabelled as least concern. The resulting dataset included 9,143 species, with 7,546 categorised as least concern (LC), 664 as near threatened, 523 as vulnerable (VU), 269 as endangered (EN), and 141 as critically endangered (CR).

To perform these analyses we first calculated the observed mean distance to centroid of all observations in the focal dataset to obtain a measure of current colour pattern diversity. We then sequentially removed species classified from the most to least threatened IUCN categories (CR>EN>VU>NT), calculating observed mean centroid distance scores for the remaining species in each case. We then used null models to test whether the observed changes in mean centroid distance after species removal was significantly different to that expected by chance, given an equivalent reduction in species richness. To do this, we sequentially removed the same number of species as present in each lost IUCN category randomly from the total species pool and measured the mean distance to centroid of the resultant null assemblage. We repeated this procedure 2,000 times and used the resulting distribution of null mean centroid distance scores to calculate both unstandardised [observed – mean(null)] and standardised [observed – mean(null)/sd(null)]. A standardised effect sizes (SES) score of >|1.96| indicates that the observed value falls outside two standard deviations of the null mean, equivalent to a *p*-value of 0.05. In any case, we calculated a *p*-value for each effect relative to null expectations using the formula described in Cooney *et al.* (*54*).

We also repeated this procedure for different subsets of our plumage embedding datasets, including (i) sex, taxon and biome-specific subsets (see below), (ii) using alternative trait diversity metrics (primary analyses only) and (iii) for related response variables including adjacency analysis parameters and species’ aesthetic attractiveness scores (see below).

### Species-level analyses

To test species-level associations between variables while controlling for sources of non-independence between observations, we used Bayesian phylogenetic mixed models (BPMMs) implemented in the R package MCMCglmm (v2.35) (*55*, *56*). BPMMs included a standard phylogenetic random effects term consisting of an inverse phylogenetic covariance matrix based on a maximum clade credibility tree for all Jetz *et al.* taxa (Hackett backbone). Where appropriate, models also included ‘species’ and ‘sex’ random effect terms to account for repeated observations per species and sex-specific variance components. In all cases, models were run for 55,000 iterations (sampled every 25th iteration) with a 5,000-iteration burn-in and using standard non-informative priors (e.g. list(R = list(V = 1, nu = 0.002), G = list(G1 = list(V = 1, nu = 0.002)))).

### Geographic analyses

We based our geographic analyses on bird species’ distribution maps produced by BirdLife International (http://datazone.birdlife.org/) previously compiled by Cooney *et al.* (*11*). Briefly, we resolved taxonomic differences between the BirdLife and Jetz *et al.* taxonomies as far as possible, manually editing (that is, combining or splitting) distribution maps for BirdLife taxa where necessary. We focused on species’ breeding geographic ranges only (seasonality, 1 or 2) and regions where species are known to be native or reintroduced (origin, 1 or 2) and extant or probably extant (presence, 1 or 2). Species’ range maps were then projected onto an equal-area grid (Behrmann projection) at 0.5° resolution (∼50 km at the equator) and clipped to terrestrial grid cells only. We then intersected this species-level presence-absence matrix with similarly-projected spatial polygons for Earth’s 14 terrestrial biome types (*57*) (available at: https://ecoregions.appspot.com/) to generate species assemblage lists for each biome. Simulated extinction analyses (see above) were then performed separately for each biome-specific assemblage to explore biome-level spatial variation in avian plumage diversity and the degree to which it is threatened by species extinctions. Spatial analyses were performed in R principally using the packages ‘sf’ (v1.0-14) (*58*) and ‘terra’ (v1.7.55) (https://github.com/rspatial/terra).

### Aesthetic attractiveness

Scores of bird species’ visual aesthetic attractiveness to humans were sourced from the ‘iratebirds’ citizen-science database (*23*). This dataset provides average attractiveness ratings for 7060 monomorphic species, as well as male and female-specific ratings for many dichromatic species (2963 and 1164, respectively). We integrated this dataset with our own using the taxonomic crosswalk to Jetz *et al.* species names produced by Santangelli et al. (*6*).

---

**Fig. S1.**
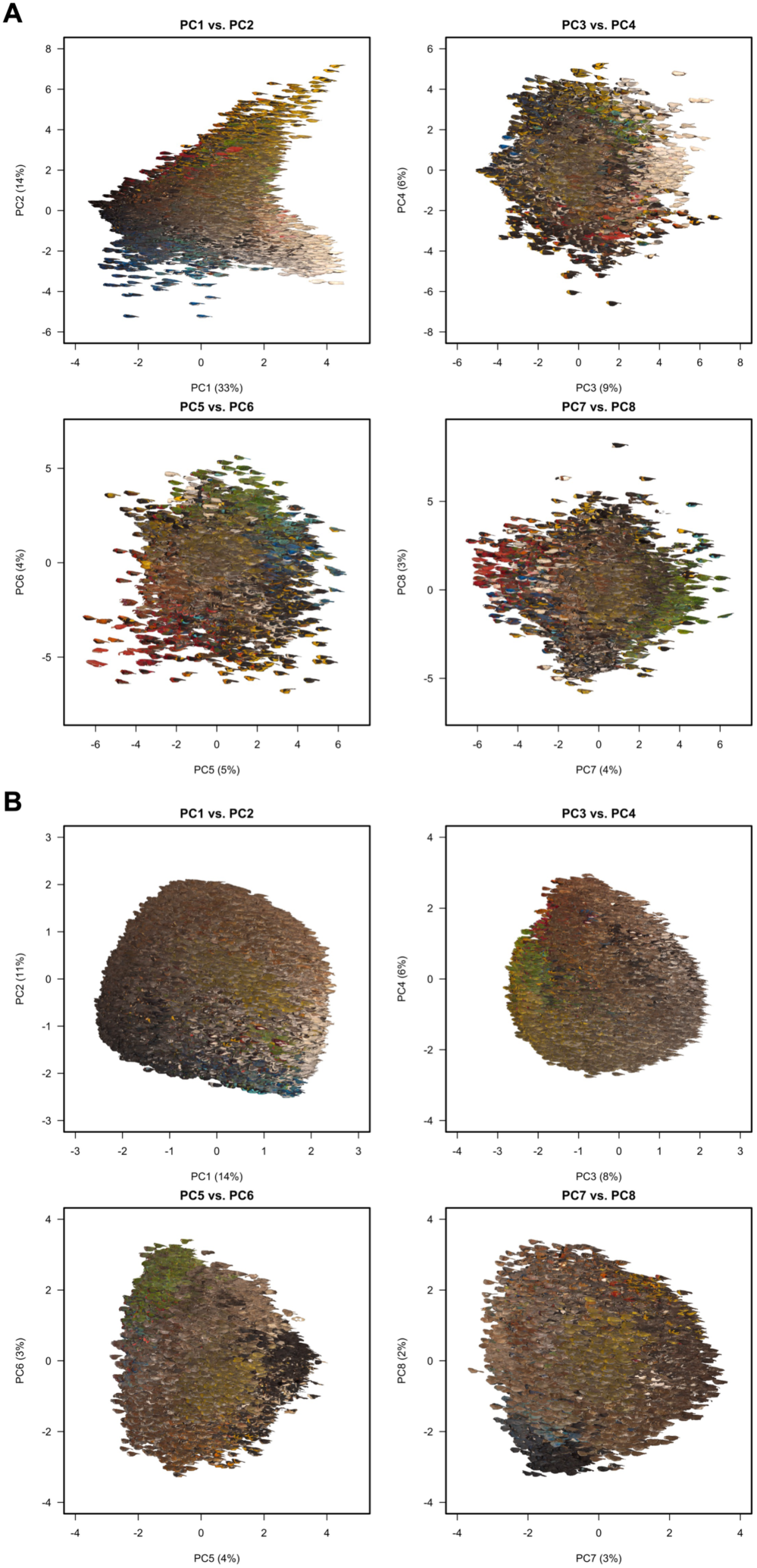
The first eight principal component (PC) axes of avian colour pattern trait space based on patch (A) and whole-plumage deep learning embeddings (B). Specimens are represented by their lateral view images. PC scores have been re-scaled (z-transformed) for clarity and axis labels give information on the variance explained by each PC. The total variance explained by the eight plotted PCs is 78% and 51% for the patch and deep learning datasets, respectively.

**Fig. S2.**
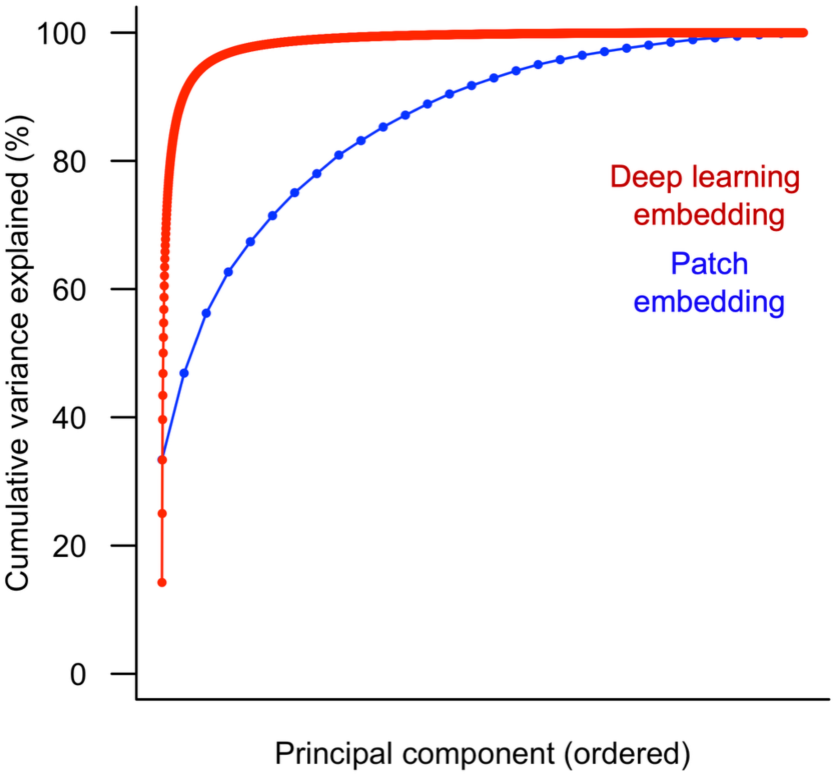
Cumulative variance explained by principal component (PC) axes of avian colour pattern trait spaces. Plotted results are based on patch (30-D) and whole-plumage deep learning embeddings (3072-D).

**Fig. S3.**
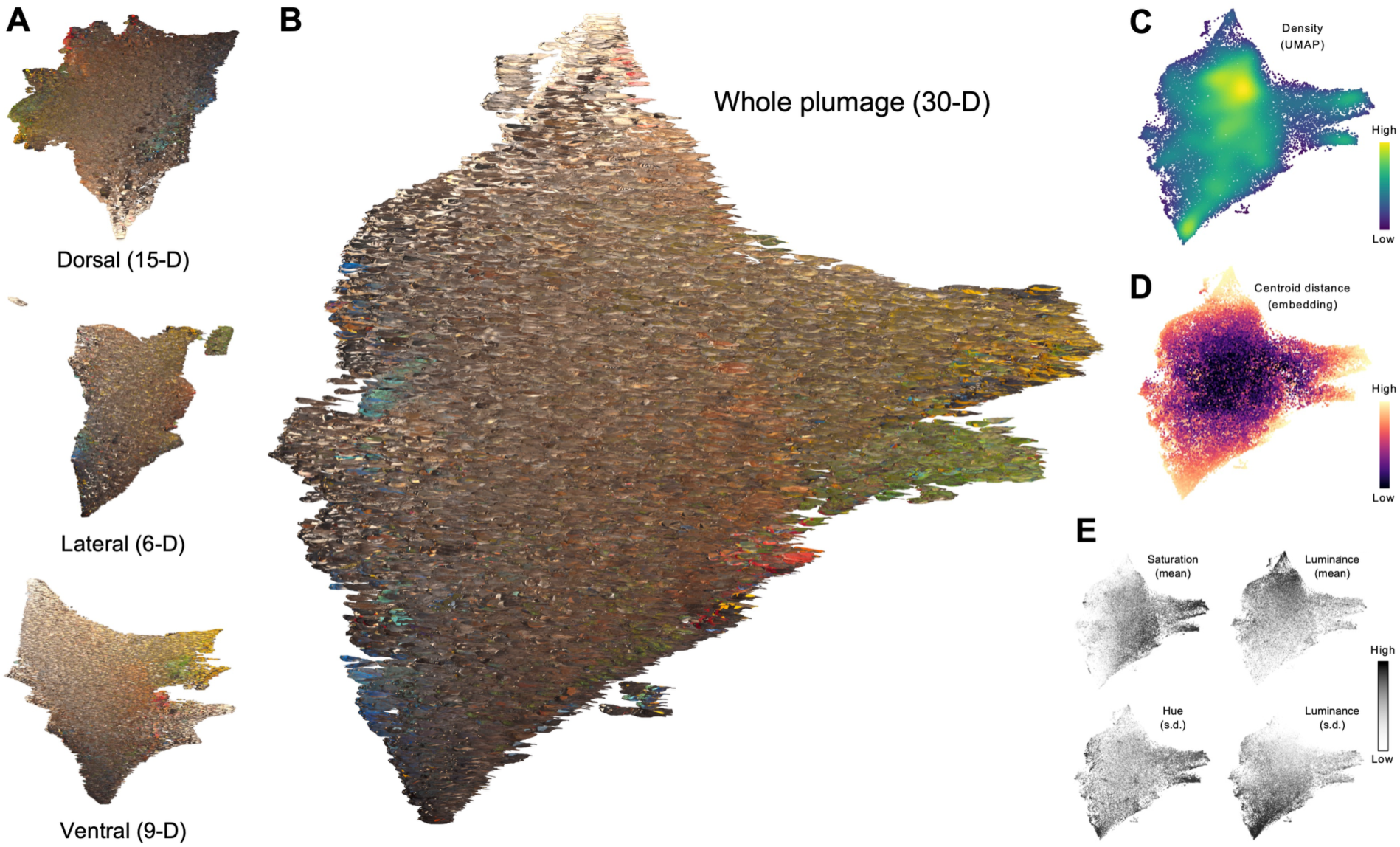
The extant avian colourscape based on patch embeddings. 2-dimensional Uniform Manifold Approximation and Projection (UMAP) visualisations of viewing angle-specific (A) and combined (B) deep learning plumage trait spaces. Note: the whole-plumage projection (B) is represented using lateral-view specimen thumbnails only. Versions of the whole-plumage projections are shown coloured according to point density in UMAP space (C), centroid distance in full embedding trait space (D) and by selected colour pattern parameter values derived from adjacency analysis (see Materials and Methods).

**Fig. S4.**
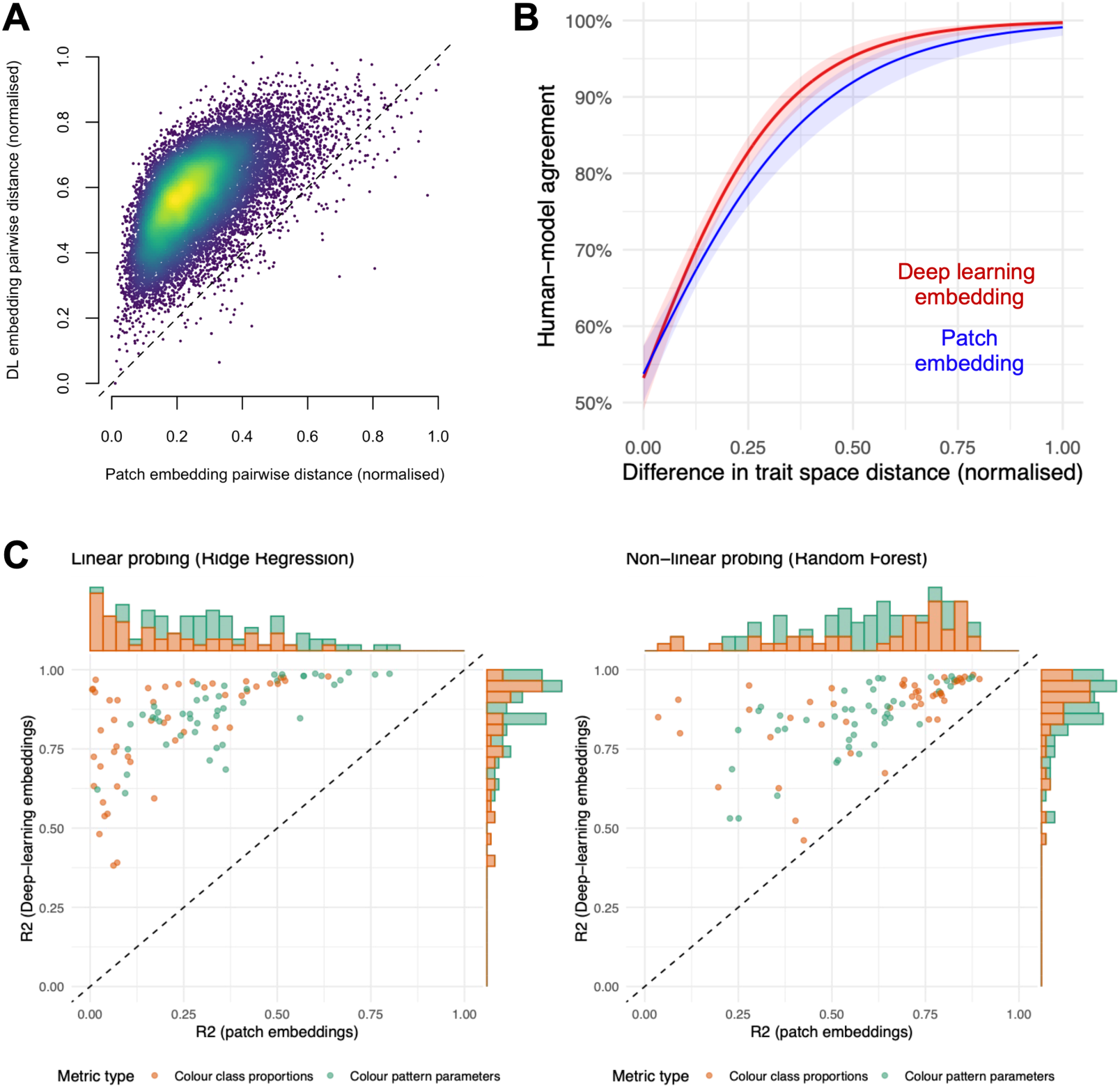
Trait space similarity, structure and interpretability. (A) The relationship between pairwise distance estimates based on patch and deep learning trait spaces for a random sample of 5,000 datapoints. Point colour/shading indicates point density (low = blue, high = yellow). (B) The fitted relationship (plus confidence interval, shaded area) for the association between human-model agreement and the magnitude of the (normalised) distance difference in embedding space. (C) Scatterplots with axis histograms showing the distribution and relationship between the proportion of variance explained (*R*^2^) by linear (left panel) and non-linear (right panel) models probing the performance of patch and deep learning trait spaces axes for explaining variation in colour class proportions (see Table S14) and colour pattern parameters (see Table S15) derived from adjacency analysis.

**Fig. S5.**
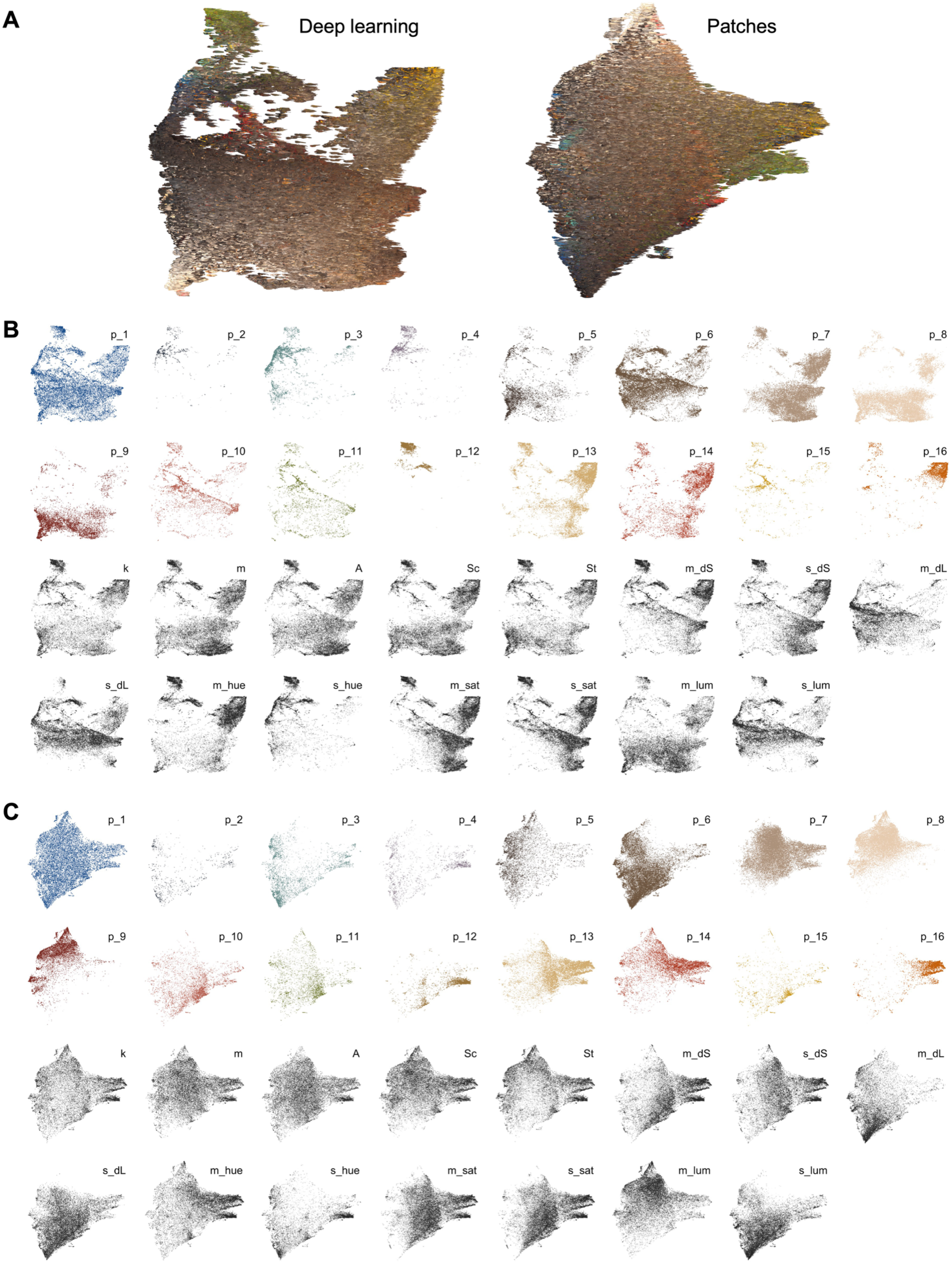
The distribution of colour pattern parameter values derived from adjacency analysis across deep learning and patch embedding spaces. (A) UMAP plots of the two embedding spaces (visualised with lateral view specimen images) for reference. (B) and (C) show the distribution of colour classes (p_*i*) and colour pattern parameters with reference to the deep learning and patch embeddings, respectively. Points in (B) and (C) are coloured according to whether the overall parameter value, averaged across viewing angles, is less than (white) or greater than (coloured) the mean parameter value across the dataset. Parameter descriptions can be found in Tables S14 and S15.

**Fig. S6.**
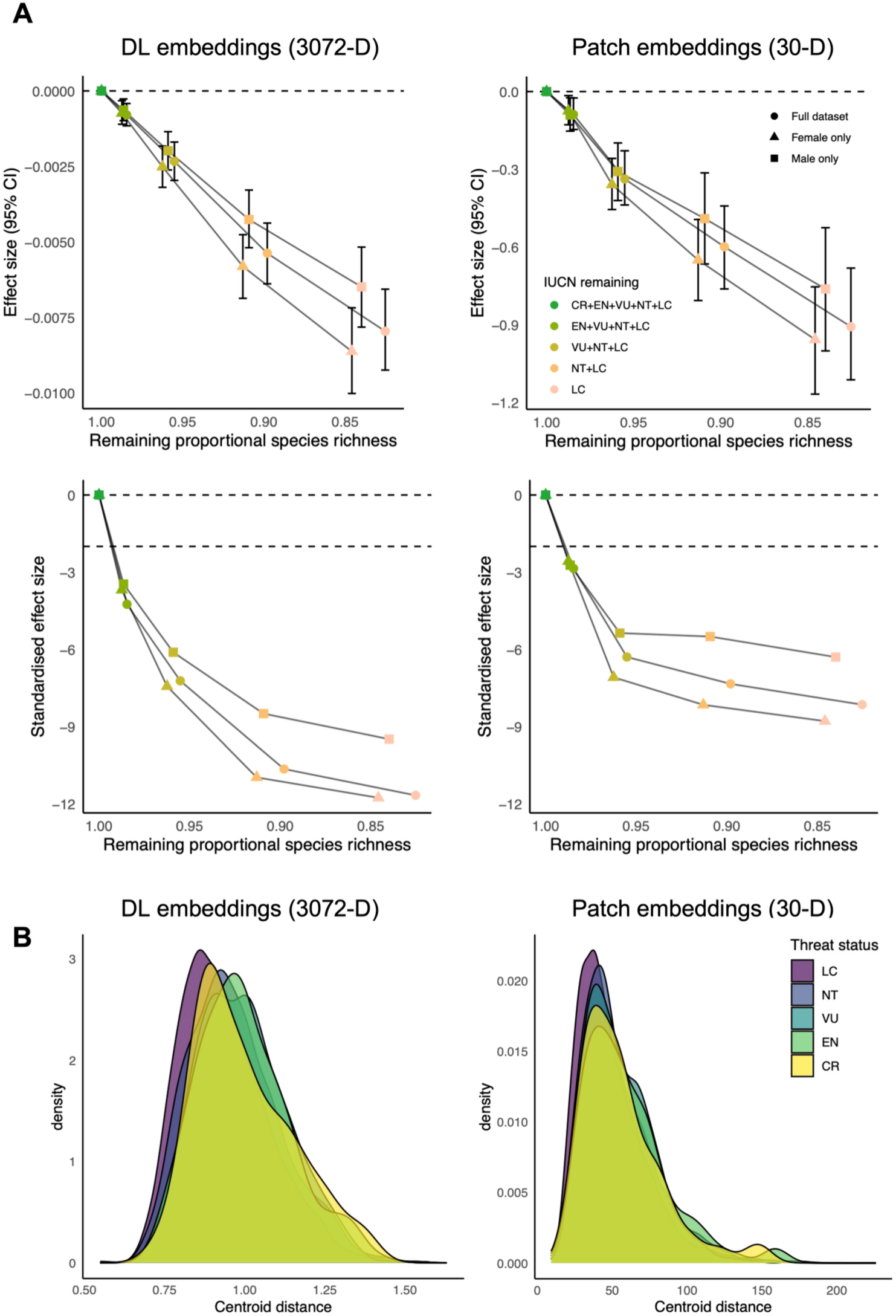
Simulated extinction results using alternative trait spaces (A) and the distribution of species centroid distance scores with respect to IUCN threat category (B). Error bars in the top plot row denotes 95% confidence interval (CI) of the parameter estimate. Note: the top left plot of panel A replicates the plot shown in Figure 2A.

**Fig. S7.**
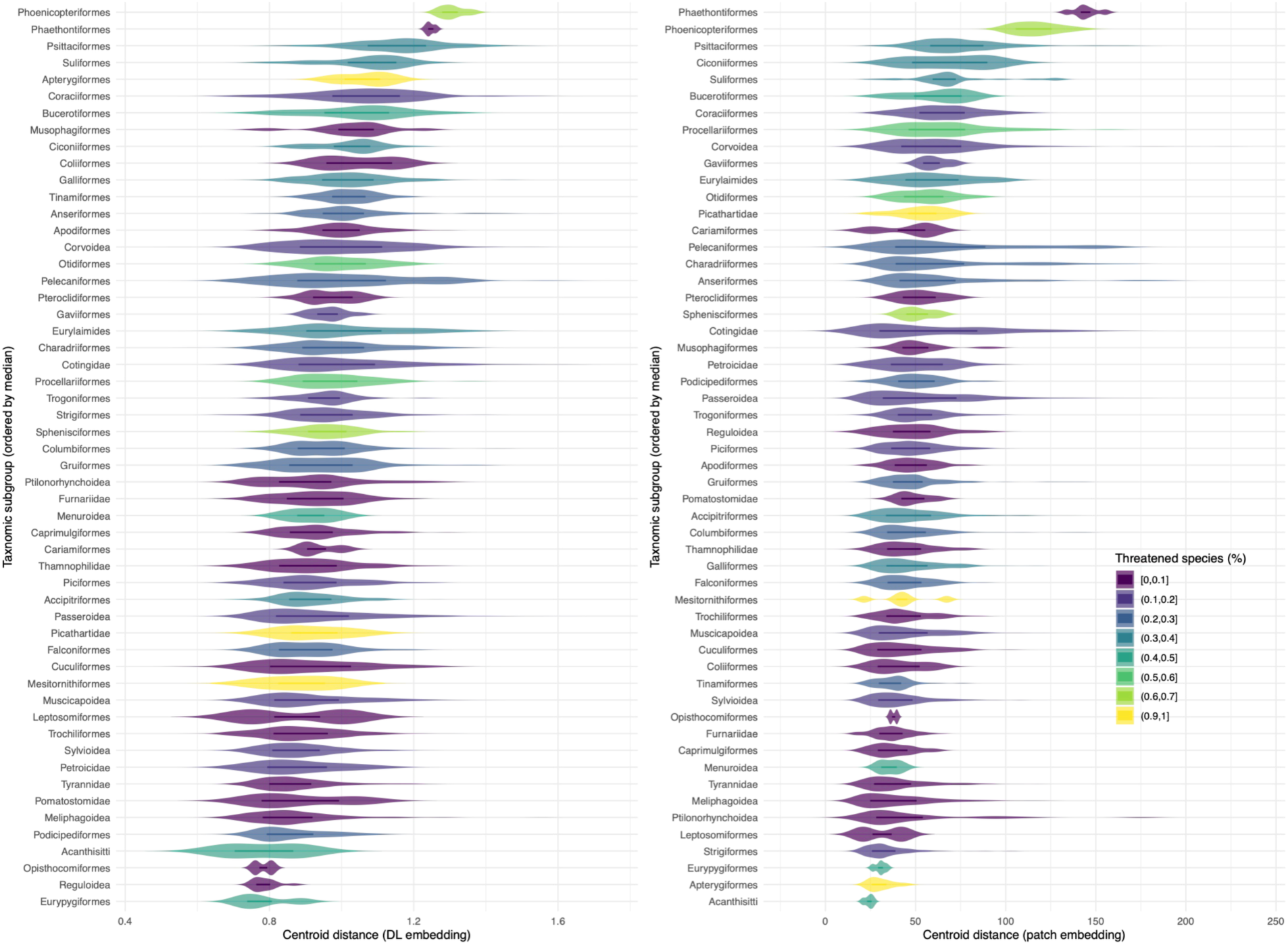
Plumage centroid distance scores with respect to taxonomic group based on whole-plumage deep learning embeddings (left) and patch embeddings (right). Violins denote the probability density distribution, and the internal boxplot displays the distribution median (center line) and interquartile range (box).

**Fig. S8.**
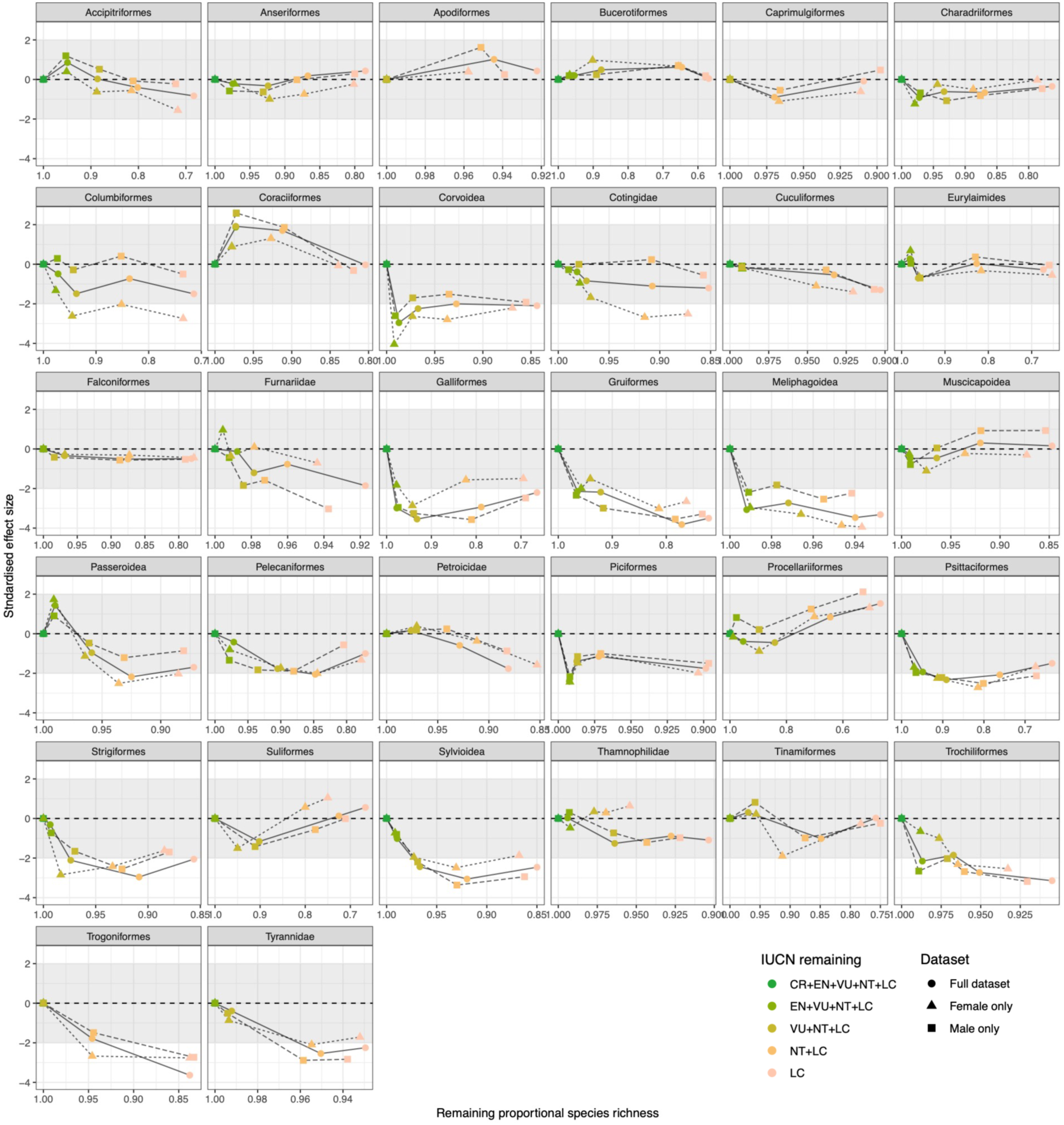
Simulated extinction results separately for major avian taxonomic groups. Only results based on the full deep learning trait space are shown. For results based on other data subsets see Table S7.

**Fig. S9.**
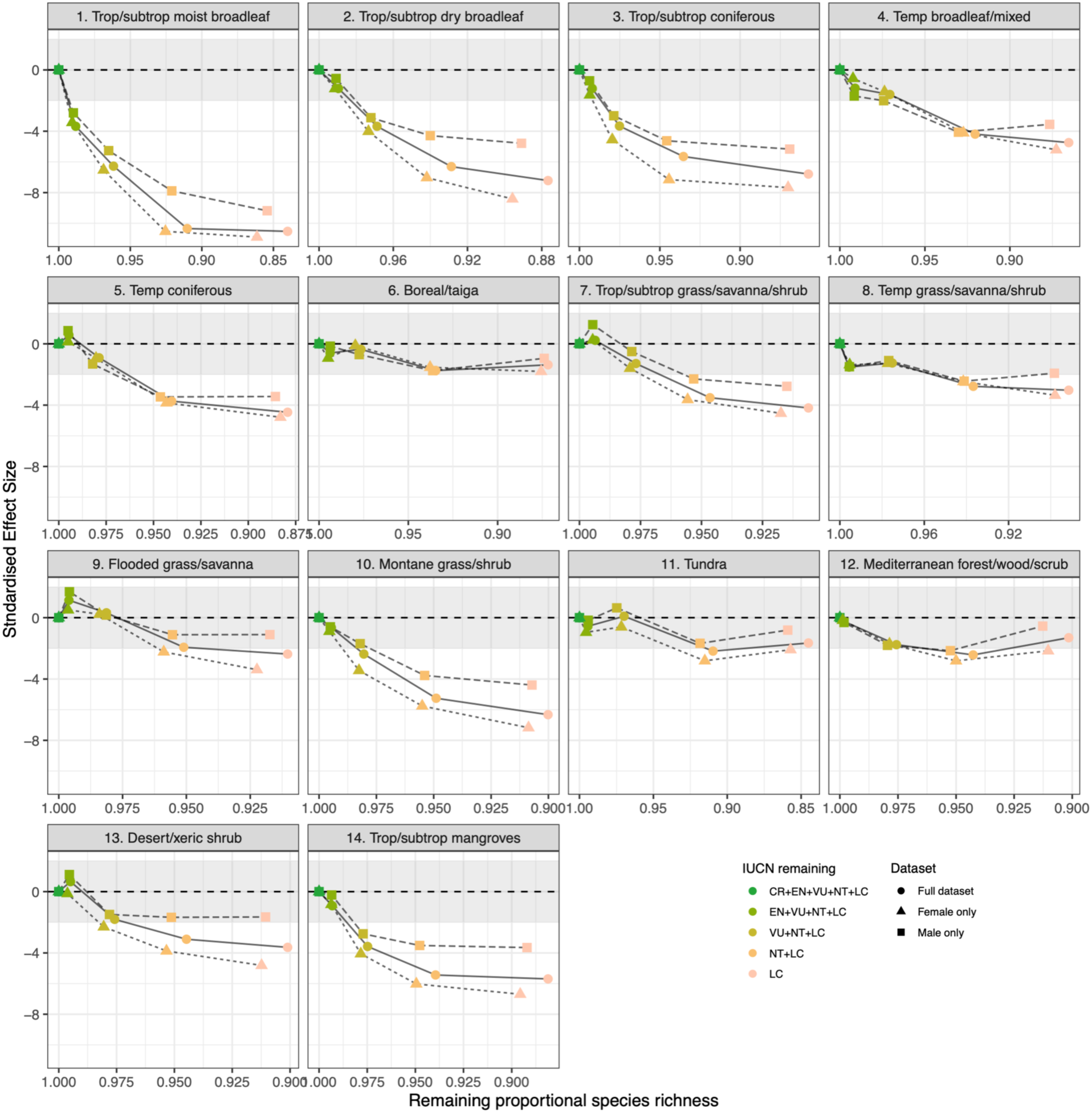
Simulated extinction results separately for biome-specific avian assemblages. Only results based on the full deep learning trait space are shown. For results based on other data subsets see Table S8.

**Fig. S10.**
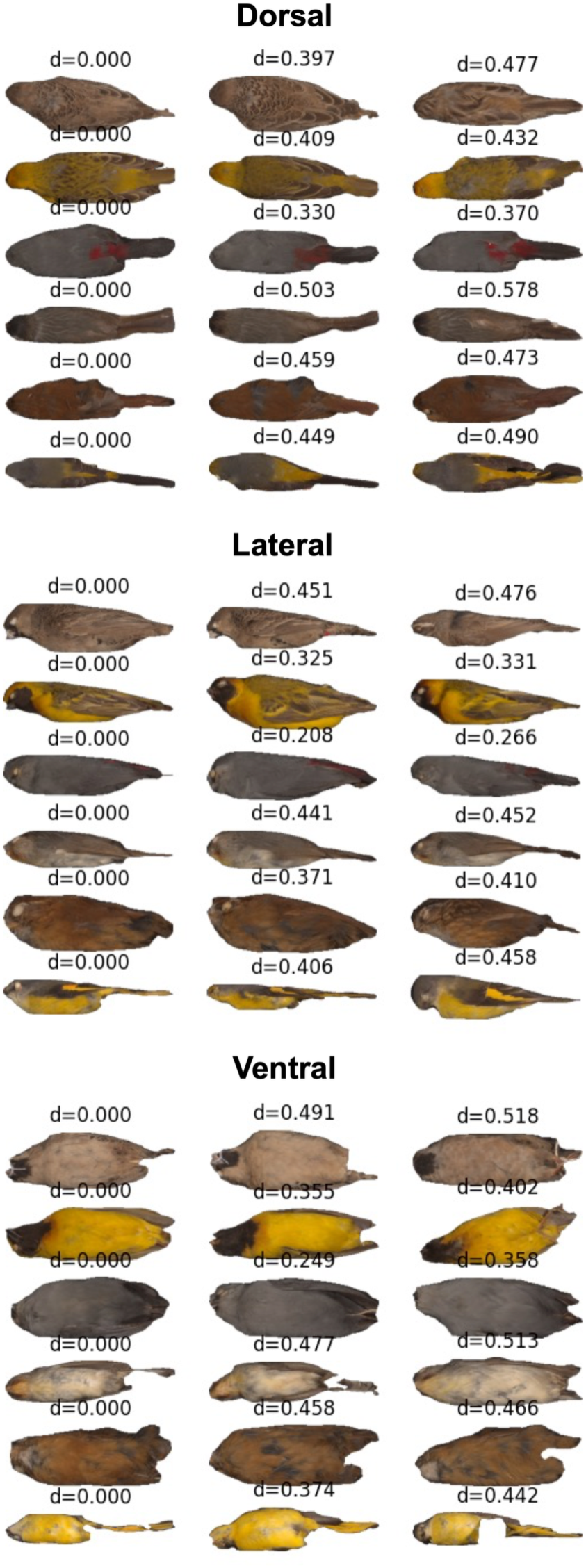
Randomly chosen examples (left column) and their nearest neighbours (middle and right columns) in the whole-plumage deep learning embedding space (SimCLR, ViT backbone). The values of *d* give the Euclidean distance between the query and the two nearest neighbours.

**Fig. S11.**
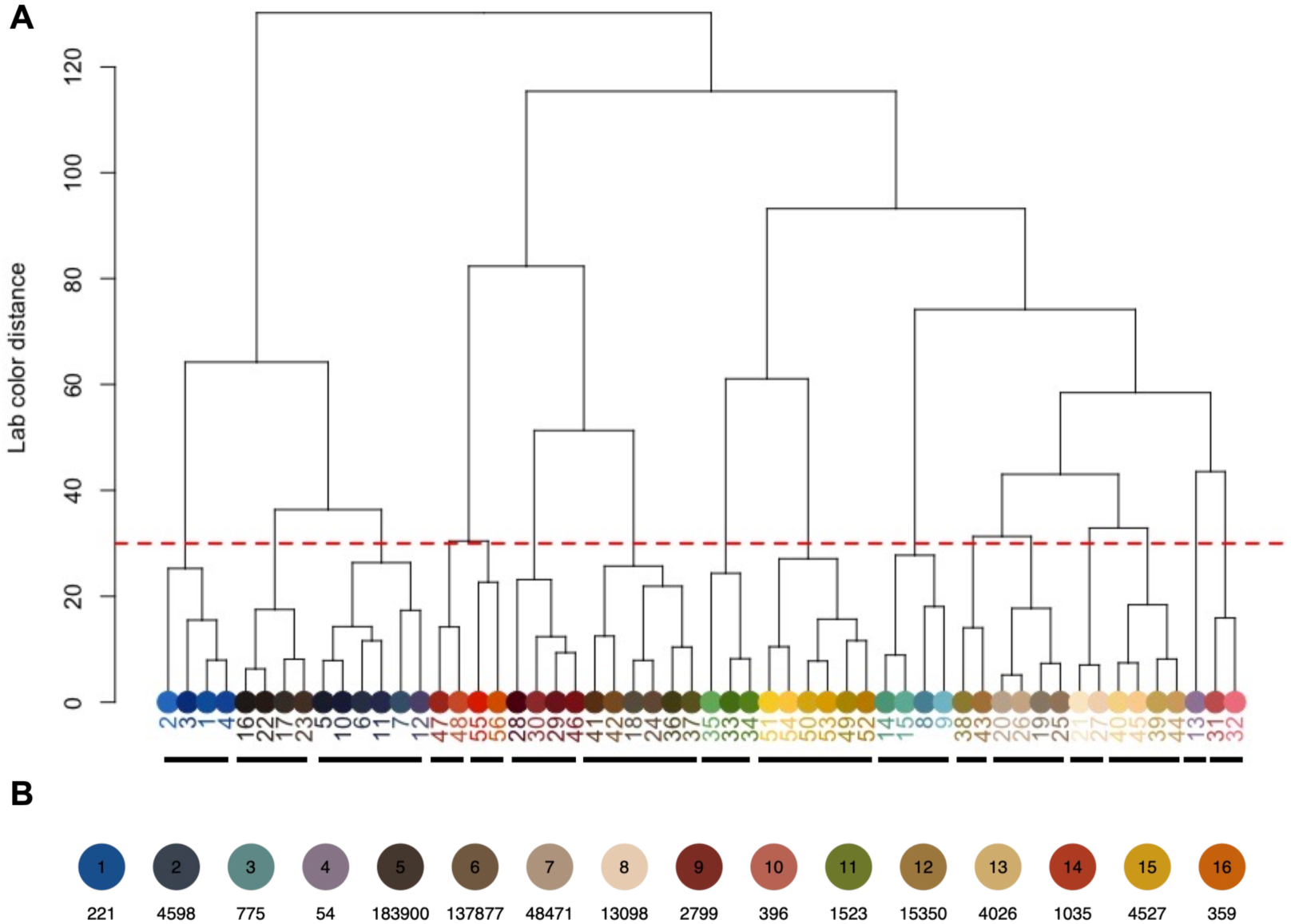
Construction of a universal set of colour classes for zone map-based adjacency analysis. See The red dashed line in (A) equates to a Lab colour distance cut-off of 30 units. Centres separated by colour distance scores of <30 are grouped to generate the final set of 16 colour classes shown in (B). Numbers beneath the coloured circles in (B) denote the number of patch observations falling into each colour class. See Table S14 for colour center RGB values and indicative names.

## Notes

### Competing Interest Statement

The authors have declared no competing interest.

## References and Notes

1. T. J. Matthews, K. A. Triantis, J. P. Wayman, T. E. Martin, J. P. Hume, P. Cardoso, S. Faurby, C. D. Mendenhall, P. Dufour, F. Rigal, R. Cooke, R. J. Whittaker, A. L. Pigot, C. Thébaud, M. W. Jørgensen, E. Benavides, F. C. Soares, W. Ulrich, Y. Kubota, J. P. Sadler, J. A. Tobias, F. Sayol, The global loss of avian functional and phylogenetic diversity from anthropogenic extinctions. Science 386, 55–60 (2024).

2. E. C. Hughes, D. P. Edwards, G. H. Thomas, The homogenization of avian morphological and phylogenetic diversity under the global extinction crisis. Current Biology 32, 3830–3837.e3 (2022).

3. J. R. Ali, B. W. Blonder, A. L. Pigot, J. A. Tobias, Bird extinctions threaten to cause disproportionate reductions of functional diversity and uniqueness. Functional Ecology 37, 162–175 (2023).

4. R. A. Graves, S. M. Pearson, M. G. Turner, Species richness alone does not predict cultural ecosystem service value. Proceedings of the National Academy of Sciences 114, 3774–3779 (2017).

5. D. R. Bellwood, C. R. Hemingson, S. B. Tebbett, Subconscious Biases in Coral Reef Fish Studies. BioScience 70, 621–627 (2020).

6. A. Santangeli, A. Haukka, W. Morris, S. Arkkila, K. Delhey, B. Kempenaers, M. Valcu, J. Dale, A. Lehikoinen, S. Mammola, What drives our aesthetic attraction to birds? npj biodivers 2, 1–7 (2023).

7. J. C. Fisher, M. Dallimer, K. N. Irvine, S. G. Aizlewood, G. E. Austen, R. D. Fish, P. M. King, Z. G. Davies, Human well-being responses to species’ traits. Nat Sustain 6, 1219–1227 (2023).

8. I. C. Cuthill, W. L. Allen, K. Arbuckle, B. Caspers, G. Chaplin, M. E. Hauber, G. E. Hill, N. G. Jablonski, C. D. Jiggins, A. Kelber, J. Mappes, J. Marshall, R. Merrill, D. Osorio, R. Prum, N. W. Roberts, A. Roulin, H. M. Rowland, T. N. Sherratt, J. Skelhorn, M. P. Speed, M. Stevens, M. C. Stoddard, D. Stuart-Fox, L. Talas, E. Tibbetts, T. Caro, The biology of color. Science 357, eaan0221 (2017).

9. T. Caro, M. C. Stoddard, D. Stuart-Fox, Animal coloration research: why it matters. Philosophical Transactions of the Royal Society B: Biological Sciences 372, 20160333 (2017).

10. M. Koneru, T. Caro, Animal Coloration in the Anthropocene. Front. Ecol. Evol. 10 (2022).

11. C. R. Cooney, Y. He, Z. K. Varley, L. O. Nouri, C. J. A. Moody, M. D. Jardine, A. Liker, T. Székely, G. H. Thomas, Latitudinal gradients in avian colourfulness. Nat Ecol Evol 6, 622–629 (2022).

12. R. A. Senior, B. F. Oliveira, J. Dale, B. R. Scheffers, Wildlife trade targets colorful birds and threatens the aesthetic value of nature. Current Biology 32, 4299–4305.e4 (2022).

13. Y. He, C. R. Cooney, S. Maddock, G. H. Thomas, PhenoLearn: a user-friendly toolkit for image annotation and deep learning-based phenotyping for biological datasets. *j. evol*. Biol. 38, 1152–1162 (2025).

14. Y. He, C. R. Cooney, S. Maddock, G. H. Thomas, Using pose estimation to identify regions and points on natural history specimens. PLOS Computational Biology 19, e1010933 (2023).

15. T. Chen, S. Kornblith, M. Norouzi, G. Hinton, A Simple Framework for Contrastive Learning of Visual Representations. arXiv arXiv:2002.05709 [Preprint] (2020). 10.48550/arXiv.2002.05709.

16. A. Puissant, A. Chotard, F. L. Condamine, V. Llaurens, Convergence in sympatric swallowtail butterflies reveals ecological interactions as a key driver of worldwide trait diversification. Proceedings of the National Academy of Sciences 120, e2303060120 (2023).

17. B. R. Scheffers, B. F. Oliveira, I. Lamb, D. P. Edwards, Global wildlife trade across the tree of life. Science 366, 71–76 (2019).

18. A. Haukka, J. Jürgens, J. Staerk, A. Lehikoinen, S. Bruslund, A. Santangeli, Aesthetic values predict bird trade, but the association varies across product types and trade regions. Biological Conservation 313, 111572 (2026).

19. J. Langlois, F. Guilhaumon, F. Baletaud, N. Casajus, C. D. A. Braga, V. Fleuré, M. Kulbicki, N. Loiseau, D. Mouillot, J. P. Renoult, A. Stahl, R. D. S. Smith, A.-S. Tribot, N. Mouquet, The aesthetic value of reef fishes is globally mismatched to their conservation priorities. PLOS Biology 20, e3001640 (2022).

20. E. Laliberté, P. Legendre, A distance-based framework for measuring functional diversity from multiple traits. Ecology 91, 299–305 (2010).

21. J. A. Endler, A framework for analysing colour pattern geometry: adjacent colours. Biol J Linn Soc 107, 233–253 (2012).

22. A. S. Tribot, J. Deter, T. Claverie, F. Guillhaumon, S. Villéger, N. Mouquet, Species diversity and composition drive the aesthetic value of coral reef fish assemblages. Biology Letters 15, 20190703 (2019).

23. A. Haukka, A. Lehikoinen, S. Mammola, W. Morris, A. Santangeli, The iratebirds Citizen Science Project: a Dataset on Birds’ Visual Aesthetic Attractiveness to Humans. Sci Data 10, 297 (2023).

24. 24. IUCN, IUCN Red List Categories and Criteria: Version 3.1 (IUCN Species Survival Commission, Gland, Switzerland and Cambridge, UK, ed. 2nd, 2012; https://portals.iucn.org/library/node/10315).

25. E. Landová, P. Poláková, S. Rádlová, M. Janovcová, M. Bobek, D. Frynta, Beauty ranking of mammalian species kept in the Prague Zoo: does beauty of animals increase the respondents’ willingness to protect them? Sci Nat 105, 69 (2018).

26. J. Marešová, D. Frynta, Noah’s Ark is full of common species attractive to humans: The case of boid snakes in zoos. Ecological Economics 64, 554–558 (2008).

27. D. Frynta, S. Lišková, S. Bültmann, H. Burda, Being Attractive Brings Advantages: The Case of Parrot Species in Captivity. PLOS ONE 5, e12568 (2010).

28. B. Lipták, A. Kouba, K. Zorić, L. Salvaras, P. Prokop, M. Paunović, The Attractiveness of Freshwater Species Correlates Positively With Conservation Support. Anthrozoös 36, 971–984 (2023).

29. W. Jetz, G. H. Thomas, J. B. Joy, K. Hartmann, A. O. Mooers, The global diversity of birds in space and time. Nature 491, 444–448 (2012).

30. C. R. Cooney, Z. K. Varley, L. O. Nouri, C. J. A. Moody, M. D. Jardine, G. H. Thomas, Sexual selection predicts the rate and direction of colour divergence in a large avian radiation. Nature Communications 10, 1773 (2019).

31. J. Troscianko, M. Stevens, Image calibration and analysis toolbox – a free software suite for objectively measuring reflectance, colour and pattern. Methods in Ecology and Evolution 6, 1320–1331 (2015).

32. R. Maia, D. R. Rubenstein, M. D. Shawkey, Selection, constraint, and the evolution of coloration in African starlings. Evolution 70, 1064–1079 (2016).

33. N. A. Mason, R. C. K. Bowie, Plumage patterns: Ecological functions, evolutionary origins, and advances in quantification. Auk 137, ukaa060 (2020).

34. C. R. Hemingson, P. F. Cowman, D. R. Bellwood, Analysing biological colour patterns from digital images: An introduction to the current toolbox. Ecology and Evolution 14, e11045 (2024).

35. Y. He, Z. K. Varley, L. O. Nouri, C. J. A. Moody, M. D. Jardine, S. Maddock, G. H. Thomas, C. R. Cooney, Deep learning image segmentation reveals patterns of UV reflectance evolution in passerine birds. Nat Commun 13, 5068 (2022).

36. A. Bellet, A. Habrard, M. Sebban, Metric Learning (Springer International Publishing, Cham, 2015; https://link.springer.com/10.1007/978-3-031-01572-4)*Synthesis Lectures on Artificial Intelligence and Machine Learning*.

37. M. D. Lürig, E. Di Martino, A. Porto, BioEncoder: A metric learning toolkit for comparative organismal biology. Ecology Letters 27, e14495 (2024).

38. K. He, X. Zhang, S. Ren, J. Sun, Deep Residual Learning for Image Recognition. arXiv arXiv:1512.03385 [Preprint] (2015). 10.48550/arXiv.1512.03385.

39. A. Dosovitskiy, L. Beyer, A. Kolesnikov, D. Weissenborn, X. Zhai, T. Unterthiner, M. Dehghani, M. Minderer, G. Heigold, S. Gelly, J. Uszkoreit, N. Houlsby, An Image is Worth 16x16 Words: Transformers for Image Recognition at Scale. arXiv arXiv:2010.11929 [Preprint] (2021). 10.48550/arXiv.2010.11929.

40. O. Siméoni, H. V. Vo, M. Seitzer, F. Baldassarre, M. Oquab, C. Jose, V. Khalidov, M. Szafraniec, S. Yi, M. Ramamonjisoa, F. Massa, D. Haziza, L. Wehrstedt, J. Wang, T. Darcet, T. Moutakanni, L. Sentana, C. Roberts, A. Vedaldi, J. Tolan, J. Brandt, C. Couprie, J. Mairal, H. Jégou, P. Labatut, P. Bojanowski, DINOv3. arXiv arXiv:2508.10104 [Preprint] (2025). 10.48550/arXiv.2508.10104.

41. E. Riba, D. Mishkin, D. Ponsa, E. Rublee, G. Bradski, Kornia: an Open Source Differentiable Computer Vision Library for PyTorch. arXiv arXiv:1910.02190 [Preprint] (2019). 10.48550/arXiv.1910.02190.

42. G. Alain, Y. Bengio, Understanding intermediate layers using linear classifier probes. arXiv arXiv:1610.01644 [Preprint] (2018). 10.48550/arXiv.1610.01644.

43. Y. Belinkov, Probing Classifiers: Promises, Shortcomings, and Advances. arXiv arXiv:2102.12452 [Preprint] (2021). 10.48550/arXiv.2102.12452.

44. H. I. Weller, A. E. Hiller, N. P. Lord, S. M. Van Belleghem, recolorize: An R package for flexible colour segmentation of biological images. Ecology Letters 27, e14378 (2024).

45. R. Maia, H. Gruson, J. A. Endler, T. E. White, R. B. O’Hara, pavo 2: new tools for the spectral and spatial analysis of colour in R. Methods in Ecology and Evolution 10, 1097–1107 (2019).

46. R. Zhang, P. Isola, A. A. Efros, E. Shechtman, O. Wang, The Unreasonable Effectiveness of Deep Features as a Perceptual Metric. arXiv arXiv:1801.03924 [Preprint] (2018). 10.48550/arXiv.1801.03924.

47. B. Hie, H. Cho, B. DeMeo, B. Bryson, B. Berger, Geometric Sketching Compactly Summarizes the Single-Cell Transcriptomic Landscape. Cell Systems 8, 483–493.e7 (2019).

48. T. Woo, X. Liang, D. A. Evans, O. Fernandez, F. Kretschmer, S. Reiter, G. Laurent, The dynamics of pattern matching in camouflaging cuttlefish. Nature 619, 122–128 (2023).

49. E. Altan, S. A. Solla, L. E. Miller, E. J. Perreault, Estimating the dimensionality of the manifold underlying multi-electrode neural recordings. PLOS Computational Biology 17, e1008591 (2021).

50. T. Guillerme, M. N. Puttick, A. E. Marcy, V. Weisbecker, Shifting spaces: Which disparity or dissimilarity measurement best summarize occupancy in multidimensional spaces? Ecology and Evolution 10, 7261–7275 (2020).

51. T. Guillerme, dispRity: A modular R package for measuring disparity. Methods in Ecology and Evolution 9, 1755–1763 (2018).

52. T. Guillerme, N. Cooper, S. L. Brusatte, K. E. Davis, A. L. Jackson, S. Gerber, A. Goswami, K. Healy, M. J. Hopkins, M. E. H. Jones, G. T. Lloyd, J. E. O’Reilly, A. Pate, M. N. Puttick, E. J. Rayfield, E. E. Saupe, E. Sherratt, G. J. Slater, V. Weisbecker, G. H. Thomas, P. C. J. Donoghue, Disparities in the analysis of morphological disparity. Biology Letters 16, 20200199 (2020).

53. J. A. Tobias, C. Sheard, A. L. Pigot, A. J. M. Devenish, J. Yang, F. Sayol, M. H. C. Neate-Clegg, N. Alioravainen, T. L. Weeks, R. A. Barber, P. A. Walkden, H. E. A. MacGregor, S. E. I. Jones, C. Vincent, A. G. Phillips, N. M. Marples, F. A. Montaño-Centellas, V. Leandro-Silva, S. Claramunt, B. Darski, B. G. Freeman, T. P. Bregman, C. R. Cooney, E. C. Hughes, E. J. R. Capp, Z. K. Varley, N. R. Friedman, H. Korntheuer, A. Corrales-Vargas, C. H. Trisos, B. C. Weeks, D. M. Hanz, T. Töpfer, G. A. Bravo, V. Remeš, L. Nowak, L. S. Carneiro, A. J. Moncada R., B. Matysioková, D. T. Baldassarre, A. Martínez-Salinas, J. D. Wolfe, P. M. Chapman, B. G. Daly, M. C. Sorensen, A. Neu, M. A. Ford, R. J. Mayhew, L. Fabio Silveira, D. J. Kelly, N. N. D. Annorbah, H. S. Pollock, A. M. Grabowska-Zhang, J. P. McEntee, J. Carlos, T. Gonzalez, C. G. Meneses, M. C. Muñoz, L. L. Powell, G. A. Jamie, T. J. Matthews, O. Johnson, G. R. R. Brito, K. Zyskowski, R. Crates, M. G. Harvey, M. Jurado Zevallos, P. A. Hosner, T. Bradfer-Lawrence, J. M. Maley, F. G. Stiles, H. S. Lima, K. L. Provost, M. Chibesa, M. Mashao, J. T. Howard, E. Mlamba, M. A. H. Chua, B. Li, M. I. Gómez, N. C. García, M. Päckert, J. Fuchs, J. R. Ali, E. P. Derryberry, M. L. Carlson, R. C. Urriza, K. E. Brzeski, D. M. Prawiradilaga, M. J. Rayner, E. T. Miller, R. C. K. Bowie, R.-M. Lafontaine, R. P. Scofield, Y. Lou, L. Somarathna, D. Lepage, M. Illif, E. L. Neuschulz, M. Templin, D. M. Dehling, J. C. Cooper, O. S. G. Pauwels, K. Analuddin, J. Fjeldså, N. Seddon, P. R. Sweet, F. A. J. DeClerck, L. N. Naka, J. D. Brawn, A. Aleixo, K. Böhning-Gaese, C. Rahbek, S. A. Fritz, G. H. Thomas, M. Schleuning, AVONET: morphological, ecological and geographical data for all birds. Ecology Letters 25, 581–597 (2022).

54. C. R. Cooney, G. H. Thomas, Heterogeneous relationships between rates of speciation and body size evolution across vertebrate clades. Nature Ecology & Evolution 5, 101–110 (2021).

55. J. D. Hadfield, S. Nakagawa, General quantitative genetic methods for comparative biology: phylogenies, taxonomies and multi-trait models for continuous and categorical characters. Journal of Evolutionary Biology 23, 494–508 (2010).

56. J. D. Hadfield, MCMC methods for multi-response generalised linear mixed models: the MCMCglmm R package. Journal of Statistical Software 33, 1–22 (2010).

57. E. Dinerstein, D. Olson, A. Joshi, C. Vynne, N. D. Burgess, E. Wikramanayake, N. Hahn, S. Palminteri, P. Hedao, R. Noss, M. Hansen, H. Locke, E. C. Ellis, B. Jones, C. V. Barber, R. Hayes, C. Kormos, V. Martin, E. Crist, W. Sechrest, L. Price, J. E. M. Baillie, D. Weeden, K. Suckling, C. Davis, N. Sizer, R. Moore, D. Thau, T. Birch, P. Potapov, S. Turubanova, A. Tyukavina, N. de Souza, L. Pintea, J. C. Brito, O. A. Llewellyn, A. G. Miller, A. Patzelt, S. A. Ghazanfar, J. Timberlake, H. Klöser, Y. Shennan-Farpón, R. Kindt, J.-P. B. Lillesø, P. van Breugel, L. Graudal, M. Voge, K. F. Al-Shammari, M. Saleem, An Ecoregion-Based Approach to Protecting Half the Terrestrial Realm. BioScience 67, 534–545 (2017).

58. E. Pebesma, Simple Features for R: Standardized Support for Spatial Vector Data. The R Journal 10, 439–446 (2018).

